# Muscle-to-Bone CX3CL1 Signaling Promotes Skeletal Repair Through CX3CR1+ Osteoprogenitors

**DOI:** 10.64898/2026.05.24.727531

**Authors:** Koji Ishikawa, Choiselle Marius, Eijiro Shimada, Puvi Nadesan, Tuyet Nguyen, Mahoko Ishikawa, Jiaul Hoque, Xinyi Ma, Makoto Nakagawa, Nicholas Allen, Koki Abe, Philip Varnadore, Tomokazu Souma, Shyni Varghese, Yasuhito Yahara, Vijitha Puviindran, Benjamin Aaron Alman

## Abstract

In the context of muscle loss, bone repair is impaired, suggesting that muscle derived signals contribute to bone regeneration. However, how muscle surrounding the injury site communicates with the bone repair niche remains unclear. Here we found that CX3CL1 expression was induced in endothelial cells in muscle surrounding a femoral bone injury site. Deletion of *Cx3cl1* impaired bone healing, demonstrating a functional role for CX3CL1 in bone repair. A CX3CL1 receptor, CX3CR1, was expressed by PDGFRα⁺ stromal progenitors and lineage tracing showed that CX3CR1 expressing osteoprogenitor lineage cells accumulated at the injury site during repair. PDGFRα⁺ stromal progenitors showed enhanced osteoblastogenesis in response to recombinant CX3CL1. In older mice, local CX3CL1 delivery increased PDGFRα⁺CX3CR1⁺ osteoprogenitor accumulation and improved bone repair. These findings identify a muscle bone signaling pathway in which endothelial CX3CL1 promotes bone repair through CX3CR1 expressing osteoprogenitors.

## Introduction

Fractures accompanied by muscle injury often exhibit delayed healing or nonunion, whereas muscle flap coverage can improve repair, demonstrating an important role for the surrounding muscle in bone regeneration^1–3^. Skeletal muscle resident mesenchymal progenitors, including fibro-adipogenic progenitor like populations, can contribute directly to the fracture callus and support chondrogenic and osteogenic repair^4–6^. Skeletal muscle also can be a secretory organ, and several muscle derived factors are linked to bone homeostasis^7–10^. However, whether injury induced muscle signals regulate local osteoprogenitor populations during bone repair remains unclear. The periosteum is a source of osteoprogenitors during bone repair and lies next to the surrounding muscle ^11–14^. As such, factors released from injured muscle could directly influence local osteoprogenitor populations in this tissue.

Chemokine signaling is an important regulator of skeletal progenitor behavior during bone repair. For example, the CCL5–CCR5 axis is required for the migration of CCR5 expressing Mx1/αSMA periosteal osteoprogenitor cells, and the CXCL12–CXCR4 pathway promotes mesenchymal progenitor homing to sites of bone injury^15,16^. However, these studies largely focused on chemokines produced within the skeletal niche. Given that skeletal muscle can function as a stress responsive secretory organ and release factors with potential skeletal effects, we sought to identify injury responsive chemokines derived from adjacent muscle to the injury site that could regulate osteoprogenitor behavior during bone repair. We therefore focused on chemokine expression in tissues containing muscle adjacent to the injury site.

CX3CL1 (fractalkine) exists in both membrane-bound and soluble forms, enabling it to function as both an adhesion molecule and a chemoattractant depending on biological context^17–19^. Although the CX3CL1–CX3CR1 axis is known to have a role in osteoblast–osteoclast coupling and bone homeostasis, recent evidence shows that CX3CL1 is robustly induced by tissue stress and released from skeletal muscle in response to exercise^20–26^. In addition, CX3CL1 has been identified as an important regulator of skeletal muscle homeostasis^27,28^. Since fracture repair slows with aging, a process often accompanied by sarcopenia, such a relationship may partly contribute to impaired repair in aging^29^. Despite this, whether CX3CL1–CX3CR1 signaling is activated in response to bone injury and whether it contributes to bone repair remain unknown.

Here, we identify CX3CL1 as an injury-induced regenerative signal derived from muscle adjacent to the injury site, and demonstrate that this response is attenuated with aging. Its receptor, CX3CR1, is expressed in PDGFRα⁺ osteoprogenitors, and CX3CL1–CX3CR1 signaling induces osteogenic activity. These findings uncover a previously unrecognized muscle-to-bone communication axis that regulates bone regeneration after injury.

## Results

### Injury induces localized CX3CL1 upregulation in muscle surrounding the injury site that is attenuated with aging

CX3CL1 expression was examined in young and older mice. Under the homeostatic conditions, mRNA expression levels in multiple tissues were largely comparable between young and older mice, although expression in bone was modestly lower in older mice (Supplementary Fig. 1a). Serum CX3CL1 levels were also comparable between young and older mice (Supplementary Fig. 1b). To determine whether CX3CL1 is regulated in response to bone injury, we created a drill hole in the femur and quantified CX3CL1 expression locally and across multiple tissues at day 3 after injury to examine the initial response (Fig. 1a and Supplementary Fig. 1c). *Cx3cl1* expression was robustly upregulated in muscle surrounding the injury site (Fig. 1b). In young mice, *Cx3cl1* expression increased approximately 21-fold in the muscle surrounding the injured bone compared with non-injured controls. The magnitude of this response was lower in older mice, with a 6-fold increase. Induction of *Cx3cl1* expression was not detected in the other organs examined during repair. We further examined whether the age-related difference in local CX3CL1 induction was reflected in the circulation, but serum CX3CL1 levels did not differ substantially between young and older mice following injury (Supplementary Fig. 1d). These results suggest that CX3CL1 function after a bone injury is primarily localized to the injury site.

**Figure 1.**
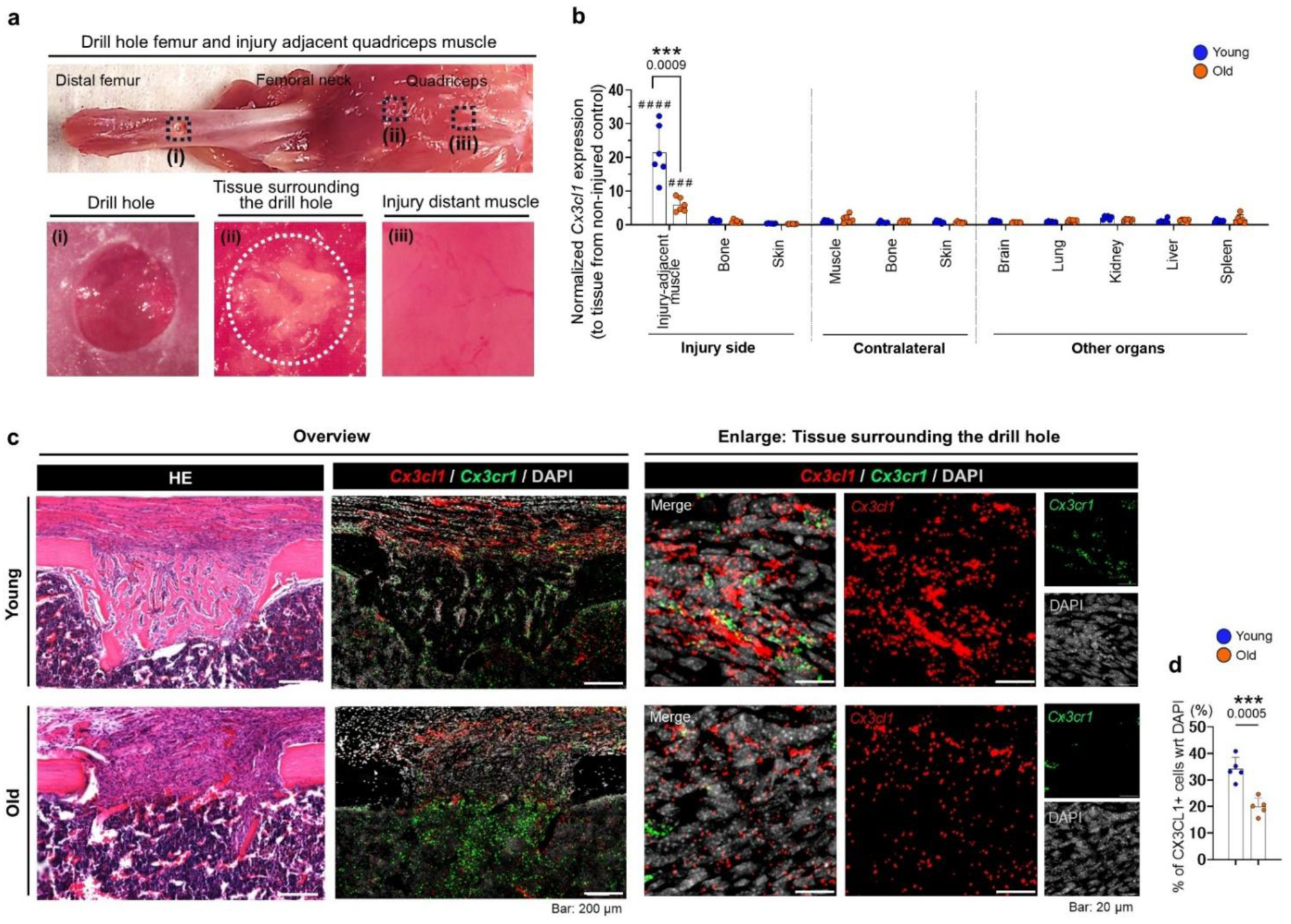
Older mice show attenuated CX3CL1 expression in muscle surrounding injured bone a, **b**, Tissue sampling and *Cx3cl1* expression at day 3 after femoral drill hole injury. a, Gross images of the drill hole site, injury adjacent quadriceps muscle, and injury distant muscle. b, qPCR analysis of *Cx3cl1* mRNA expression in injured side tissues, contralateral tissues, and other organs. Values were normalized to *Gapdh* and expressed relative to the corresponding uninjured tissue. n = 6 mice per group, 3 male and 3 female mice. ^###^ P < 0.001 and ^####^ P < 0.0001 versus uninjured mice. **c,** Representative H&E and RNAscope images for *Cx3cl1* and *Cx3cr1* in the repair region at day 7 after injury. Left, overview. Right, enlarged views of tissue surrounding the drill hole. Images are representative of n = 5 mice per group. **d,** Quantification of the percentage of *Cx3cl1*⁺ cells among DAPI⁺ cells in tissue surrounding the injured bone. n = 5 mice per group. Young mice were 3 to 4 months old, and older mice were 23 to 25 months old. Data are presented as mean ± SD. Statistical significance was determined using Student’s t test.

To define the spatial distribution of the injury induced *Cx3cl1* and the localization of a receptor, we next performed RNAscope in situ hybridization for *Cx3cl1* and *Cx3cr1* in uninjured femurs and at day 7 after drill hole injury. In uninjured femurs, *Cx3cl1* signal was minimal in cortical bone, with low level signal detected in muscle and bone marrow (Supplementary Fig. 1e). Following injury, *Cx3cl1* signal became prominent in the muscle surrounding the injured bone with a pattern suggesting expression in the vasculature (Supplementary Fig. 1f). There were more *Cx3cl1*-expressing cells in young mice than in older mice (Fig. 1c,d). In contrast, *Cx3cr1* expressing cells were primarily localized within the bone marrow at baseline but were observed surrounding newly formed bone after injury.

To define the temporal and tissue distribution of CX3CL1-expressing cells after injury, we analyzed cells isolated from muscle and bone adjacent to the injury site in young and older mice using flow cytometry at days 0, 5, and 10 after injury. CX3CL1⁺ cells were detected in muscle adjacent to the injury site, with more CX3CL1⁺ cells in young mice than in older mice (Fig. 2a,b). This difference was most pronounced at day 5 post injury, when CX3CL1⁺ cells reached peak levels in both groups. In contrast, CX3CL1 expression was minimal in bone across all time points.

**Figure 2.**
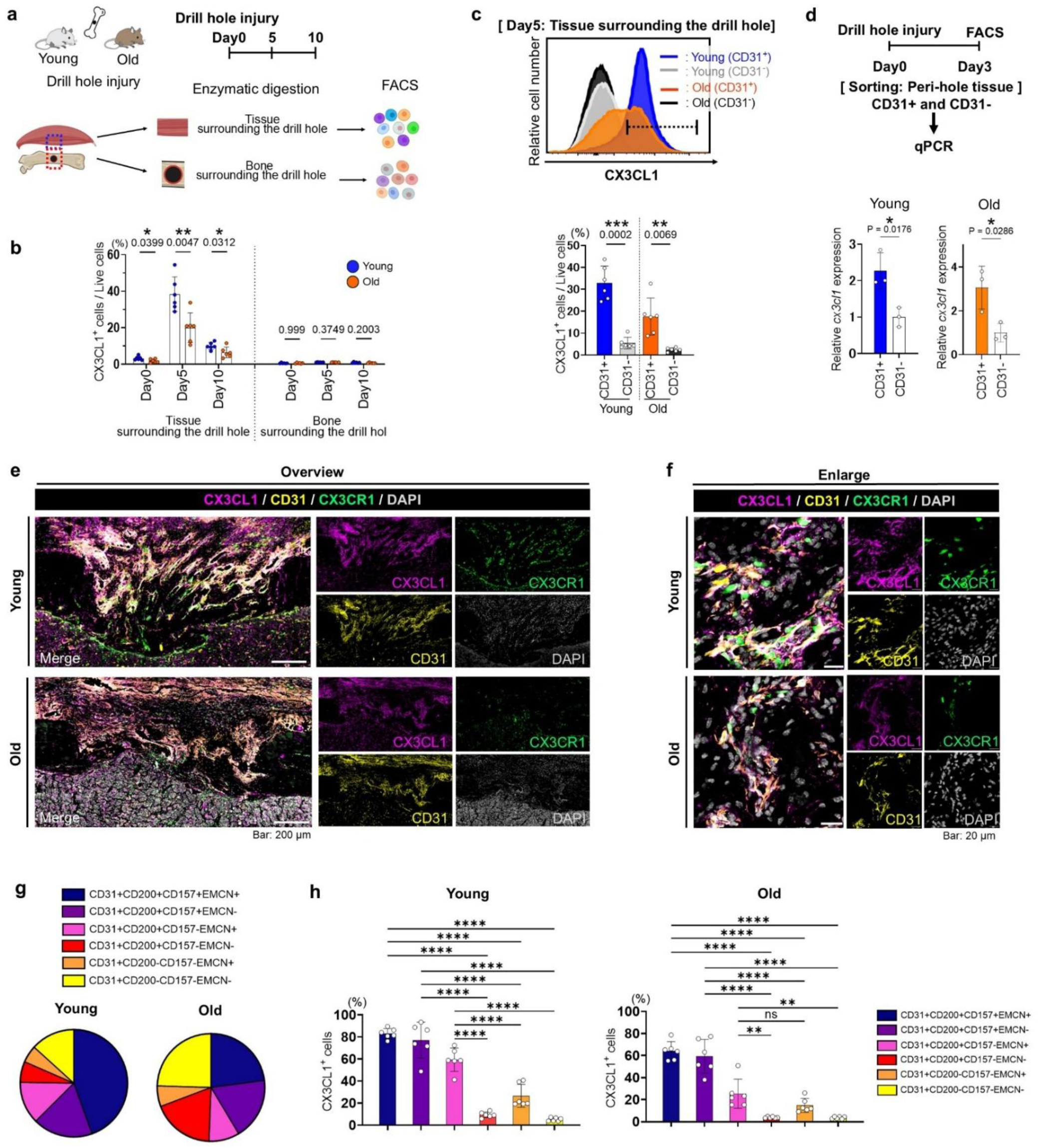
CX3CL1 is enriched in distinct endothelial subsets and attenuated in older mice. **a**, Schematic of tissue collection and flow cytometry analysis after femoral drill hole injury. **b**, Flow cytometric quantification of CX3CL1⁺ cells in muscle and bone tissue surrounding the drill hole. n = 6 mice per group, including 3 male and 3 female mice. **c,** Representative overlaid histograms and quantification of CX3CL1⁺ cells among CD31⁺ and CD31⁻ cells isolated from muscle tissue surrounding the injured bone day 5 after injury. Representative overlaid histograms were normalized to unit area. n = 6 mice per group, including 3 male and 3 female mice. **d**, Quantitative PCR analysis of *Cx3cl1* mRNA expression in sorted CD31⁺ and CD31⁻ cells isolated from muscle tissue surrounding the injured bone at day 3 after injury. n = 3 mice per group. **e, f,** Representative immunofluorescence images of the injured bone in young and older *Cx3cr1* GFP/+ mice at day 7 after injury. e, overview images. f, magnified views of drill hole area. Images are representative of n = 5 mice per group. **g**, Distribution of endothelial subsets among CD31⁺ cells isolated from muscle tissue surrounding the injured bone at day 5 after injury. n = 6 mice per group. **h**, Quantification of CX3CL1⁺ cells within each endothelial subset isolated from muscle tissue surrounding the injured bone at day 5 after injury. n = 6 mice per group. Data are presented as mean ± SD. Statistical significance was determined using Student’s t test for two group comparisons and one way ANOVA with Tukey’s multiple comparisons test for endothelial subset comparisons.

We then examined whether CX3CL1⁺ was expressed in cells expressing CD31, a marker of endothelial cells^30,31^. Flow cytometric analysis showed that CX3CL1⁺ cells were enriched in the CD31⁺ population compared with CD31⁻ cells (Fig. 2c). In both young and older mice, *Cx3cl1* expression was significantly higher in CD31⁺ cells than in CD31⁻ cells from injury adjacent muscle (Fig. 2d). To validate these findings spatially, we performed immunofluorescence staining using *Cx3cr1* GFP/+ reporter mice, in which GFP is inserted into the *Cx3cr1* locus^32^. CD31 and CX3CL1 signals largely overlapped in muscle surrounding the injury site in both young and older mice, and CX3CR1⁺ cells were positioned adjacent to these CX3CL1⁺ endothelial regions (Fig. 2e,f).

We next assessed CX3CL1 expression across CD31⁺ endothelial subsets in injury adjacent muscle, including subsets previously associated with regenerative endothelial populations (Supplementary Fig. 2a)^33–35^. At day 5 after injury, endothelial subset composition differed between young and older mice, with CD200⁺CD157⁺EMCN⁺, CD200⁺CD157⁺EMCN⁻, and CD200⁺CD157⁻EMCN⁺ subsets representing a larger fraction of CD31⁺ endothelial cells in young mice (Fig. 2g and Supplementary Fig. 2b). Notably, CX3CL1⁺ cells were enriched within these subsets, but this subset associated CX3CL1 response was reduced in older mice (Fig. 2h and Supplementary Fig. 2c). Together, these findings indicate that injury induced CX3CL1 expression is preferentially enriched within endothelial subsets associated with regenerative activity in muscle adjacent to the injury site.

### Presence of non-hematopoietic CX3CR1⁺ cells in osteoprogenitors during bone repair

Given that CX3CL1 was predominantly produced by endothelial cells in muscle surrounding the injury site, we next asked whether CX3CR1, the receptor for CX3CL1, is present within the bone repair compartment. CX3CR1 was expressed primarily in immune cells and osteoclast lineage cells^17,25,26^. Within CD45⁺ hematopoietic cells, CX3CR1 expression was most prominent in monocyte and macrophage populations, followed by neutrophils, with smaller fractions detected in dendritic cells, NK cells, and B cells. These CX3CR1⁺ hematopoietic populations were more abundant in older mice across time points (Supplementary Figs. 3a,b and 4a–i). Notably, a distinct CX3CR1⁺ population was detected within the Lin⁻CD45⁻ compartment, and this non-hematopoietic population was more abundant in young mice during the early phase of repair (Fig. 3a,b). Immunofluorescence analysis in young *Cx3cr1* GFP/+ reporter mice at day 7 after injury further confirmed the presence of CD45⁻CX3CR1 GFP⁺ cells in the drill hole repair region. These cells were enriched along bone surfaces and at the interface between bone and marrow, with fewer cells detected distant from bone injury (Fig. 3c-e). Staining for CD45 and F4/80 confirmed that a subset of CX3CR1 GFP⁺ cells in the drill hole area was neither CD45⁺ hematopoietic cells nor F4/80⁺ macrophages (Supplementary Fig. 3d,e).

**Figure 3.**
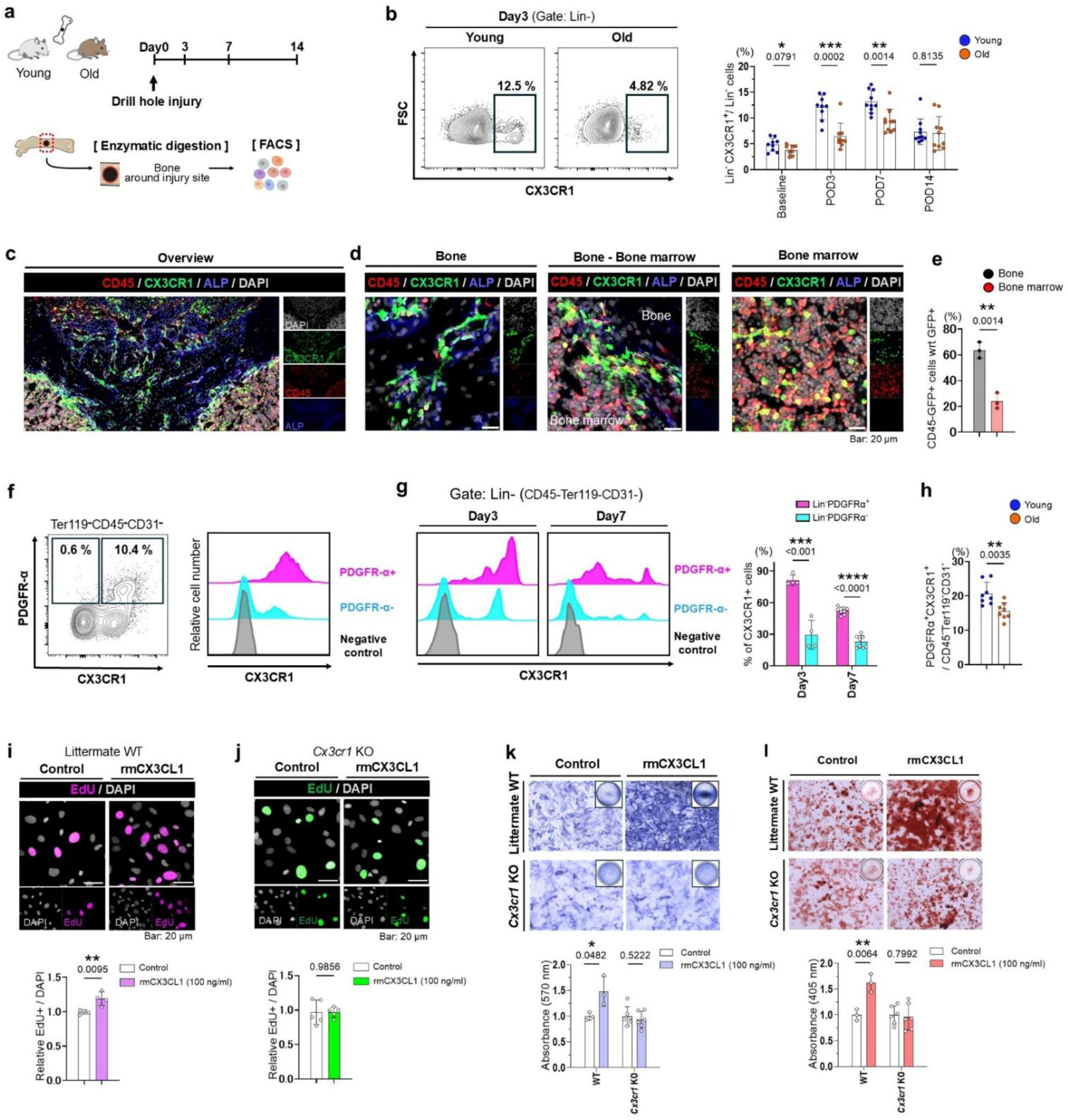
CX3CR1 identifies nonhematopoietic osteoprogenitors responsive to CX3CL1 during bone repair. **a**, Schematic of bone tissue collection and flow cytometry analysis after femoral drill hole injury. **b**, Representative flow cytometry plots and quantification of CX3CR1⁺ cells among lineage negative nonhematopoietic cells. n = 8 to 10 mice per group, including 4 to 5 males and 4 to 5 females. **c–e**, CX3CR1 GFP⁺ cells in the drill hole region of young *Cx3cr1* GFP/+ reporter mice. c, Overview of the injury region. d, Enlarged views of the injured bone, bone–marrow interface, and distant marrow region. e, Quantification of nonhematopoietic CD45⁻CX3CR1 GFP⁺ cells. n = 3 mice per group. **f,** Flow cytometric analysis of Lin⁻ nonhematopoietic cells (Ter119⁻CD45⁻CD31⁻) from uninjured young bone. Left, PDGFRα and CX3CR1 gating. Right, unit area normalized CX3CR1 histograms in PDGFRα⁺ and PDGFRα⁻ cells. Plots are representative of n = 3 mice. **g,** CX3CR1 histograms in PDGFRα⁺ and PDGFRα⁻ Lin⁻ nonhematopoietic cells at days 3 and 7 after injury. Histograms were normalized to unit area. Right, quantification of CX3CR1⁺ cells within each subset. n = 4 to 8 mice per group. **h,** Flow cytometric quantification of PDGFRα⁺CX3CR1⁺ cells among Lin⁻ nonhematopoietic cells at day 7 after injury. n = 8 mice per group, 4 male and 4 female. **i,j,** Representative EdU staining and quantification of EdU⁺ cells in CD45 depleted mesenchymal stromal cells from young littermate wild type mice from the *Cx3cr1* GFP colony (i) and *Cx3cr1* GFP/GFP mice lacking functional CX3CR1 (j). Cells were treated with vehicle or rmCX3CL1. n = 4 for i and n = 5 for j. **k,l,** ALP (k) and Alizarin Red (l) staining and quantification in CD45 depleted mesenchymal stromal cells from young littermate WT and *Cx3cr1* GFP/GFP mice cultured in osteogenic medium with vehicle or rmCX3CL1. n = 3 WT and n = 6 *Cx3cr1* GFP/GFP mice. Data are presented as mean ± SD. Statistical significance was determined using Student’s t test.

This localization pattern raised the possibility that CX3CR1 is expressed by osteoprogenitors within the bone repair niche. To test this, we examined CX3CR1 expression in PDGFRα⁺ cells within the CD45⁻Ter119⁻CD31⁻ compartment, which contains stromal progenitors that contribute to bone repair, including periosteum associated osteoprogenitors^36,37^. In young mice, CX3CR1 was enriched in PDGFRα⁺ cells relative to PDGFRα⁻ cells at baseline (Fig. 3f and Supplementary Fig. 5), and this enrichment persisted during the early phase of repair at days 3 and 7 after injury (Fig. 3g). Moreover, the PDGFRα⁺CX3CR1⁺ stromal subset was significantly more abundant in young mice than in older mice (Fig. 3h).

Given the enrichment of CX3CR1 in PDGFRα⁺ stromal cells, we next investigated whether these cells functionally respond to CX3CL1. We established mesenchymal stromal cell cultures from the CD45 depleted fraction and assessed their response to recombinant CX3CL1^38^. CX3CL1 treatment increased cell proliferation (EdU incorporation) and enhanced cell viability (Fig. 3i and Supplementary Fig. 6a). These effects were absent in cultures derived from *Cx3cr1* deficient mice (Fig. 3j and Supplementary Fig. 6b). CX3CL1 increased alkaline phosphatase activity and mineralization, whereas these osteogenic effects were not observed in *Cx3cr1* deficient cultures (Fig. 3k,l and Supplementary Fig. 6c). Thus, CX3CR1 identifies a non-hematopoietic stromal population that responds to CX3CL1 by enhancing osteogenesis.

### Spatial distribution of non-hematopoietic CX3CR1⁺ osteoprogenitors during bone repair

We next examined the spatial distribution of CX3CR1 expressing cells within the PDGFRα lineage, using *Pdgfra*-CreERT2;R26-tdTomato;*Cx3cr1*-GFP/+ mice (combining the *Pdgfra*-CreERT2, R26-tdTomato, and *Cx3cr1*-GFP reporter alleles), enabling PDGFRα lineage tracing together with GFP based identification of CX3CR1 expressing cells (Fig. 4a and Supplementary Fig. 7a). In the contralateral uninjured femur, Tomato⁺ cells were observed primarily in the periosteal region, and a substantial proportion of these Tomato⁺ cells also expressed CX3CR1 GFP (Fig. 4b). Double positive cells were also observed in several other organs (Supplementary Fig. 7b). Following drill-hole injury, a marked increase in Tomato⁺ cells was observed at the injury site by day 7, with the majority co-expressing CX3CR1-GFP (Fig. 4c,d). This pattern persisted at days 14, 21, and 28. Notably, Tomato⁺CX3CR1 GFP⁺ cells were frequently found lining newly formed bone surfaces, suggesting that PDGFRα lineage CX3CR1 expressing cells participate in new bone formation during repair.

**Figure 4.**
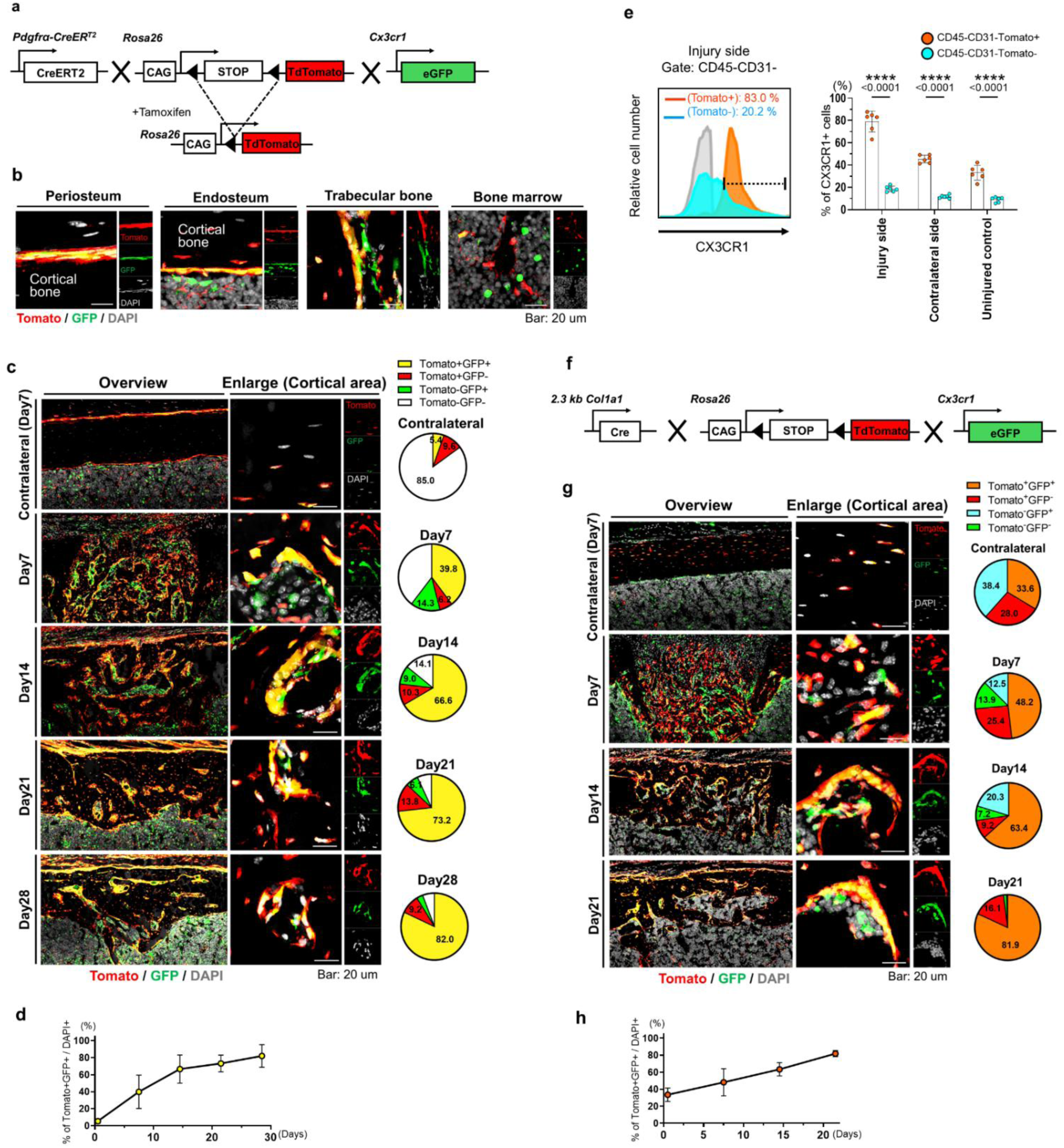
CX3CR1 expressing osteoprogenitor lineage cells localize to new bone surfaces during repair a, Genetic reporter strategy in *Pdgfra*-CreERT2; tdTomato; *Cx3cr1* GFP/+ mice. **b,** Representative fluorescence images of contralateral femoral bone from *Pdgfra*-CreERT2; tdTomato; *Cx3cr1* GFP/+ mice at day 7 after injury. Images are representative of n = 3 mice. **c,** Representative fluorescence images of the injured cortical region in *Pdgfra*-CreERT2; tdTomato; *Cx3cr1* GFP/+ mice during repair. tdTomato is shown in red, CX3CR1 GFP reporter signal in green, and DAPI in gray. Pie charts show tdTomato and GFP cell population proportions. Images are representative of n = 3 mice per time point. **d,** Quantification of tdTomato⁺GFP⁺ cells during repair. Data are mean ± 95% CI. n = 3 mice per time point. **e,** Flow cytometric analysis of CX3CR1⁺ cells in *Col1a1* lineage cells after femoral drill hole injury. Left, representative CX3CR1 histograms in CD45⁻CD31⁻tdTomato⁺ and CD45⁻CD31⁻tdTomato⁻ cells. Right, quantification of CX3CR1⁺ cells in tdTomato⁺ and tdTomato⁻ populations from injured, contralateral, and uninjured bone. n = 6 mice per group. Data are mean ± SD. Statistical significance was determined using Student’s t test. **f,** Genetic reporter strategy in Col1a1-Cre; tdTomato; *Cx3cr1* GFP/+ mice. **g**, Representative fluorescence images of the injured cortical region in *Col1a1*-Cre; tdTomato; *Cx3cr1* GFP/+ mice during repair. Pie charts show tdTomato and GFP cell population proportions. Images are representative of n = 3 mice per time point. **h**, Quantification of tdTomato⁺GFP⁺ cells during repair. Data are mean ± 95% CI. n = 3 mice per time point.

We then examined *Col1a1*-Cre; R26Tomato mice, in which osteoblast lineage cells are labeled with tdTomato. Within the CD45⁻CD31⁻Tomato⁺ population, the proportion of CX3CR1⁺ cells was markedly increased at the injury site compared with the contralateral side and uninjured mice (Fig. 4e and Supplementary Fig. 8). We next used *Col1a1*-Cre; R26-tdTomato; *Cx3cr1*-GFP/+ mice (combining the *Col1a1*-Cre, R26-tdTomato, and *Cx3cr1*-GFP reporter alleles) to examine the localization of CX3CR1 expressing cells within the osteoblast lineage during bone repair (Fig. 4f). Following injury, Tomato⁺CX3CR1-GFP⁺ cells were observed from days 7 to 21. These double positive cells were frequently observed lining newly formed bone surfaces, a finding consistent with PDGFRα lineage tracing results (Fig. 4g,h). Collectively, these findings indicate that CX3CR1 is expressed by osteogenic lineage cells that localize to sites of new bone formation during repair.

### CX3CL1 promotes bone repair by regulating CX3CR1⁺ osteoprogenitors

To examine the functional role of CX3CL1 during bone repair, we generated UBC-CreERT2;*Cx3cl1* flox/flox mice to induce global *Cx3cl1* deletion. In the absence of tamoxifen, μCT and histological analyses showed comparable bone repair between groups after drill hole injury (Supplementary Fig. 9a-e). We next induced *Cx3cl1* deletion using tamoxifen (Fig. 5a). Efficient deletion was confirmed by significantly lower serum CX3CL1 levels in deleted mice than in littermate controls (Fig. 5b). Since *Cx3cl1* is expressed in neural tissues, we performed open field testing under uninjured conditions.

**Figure 5.**
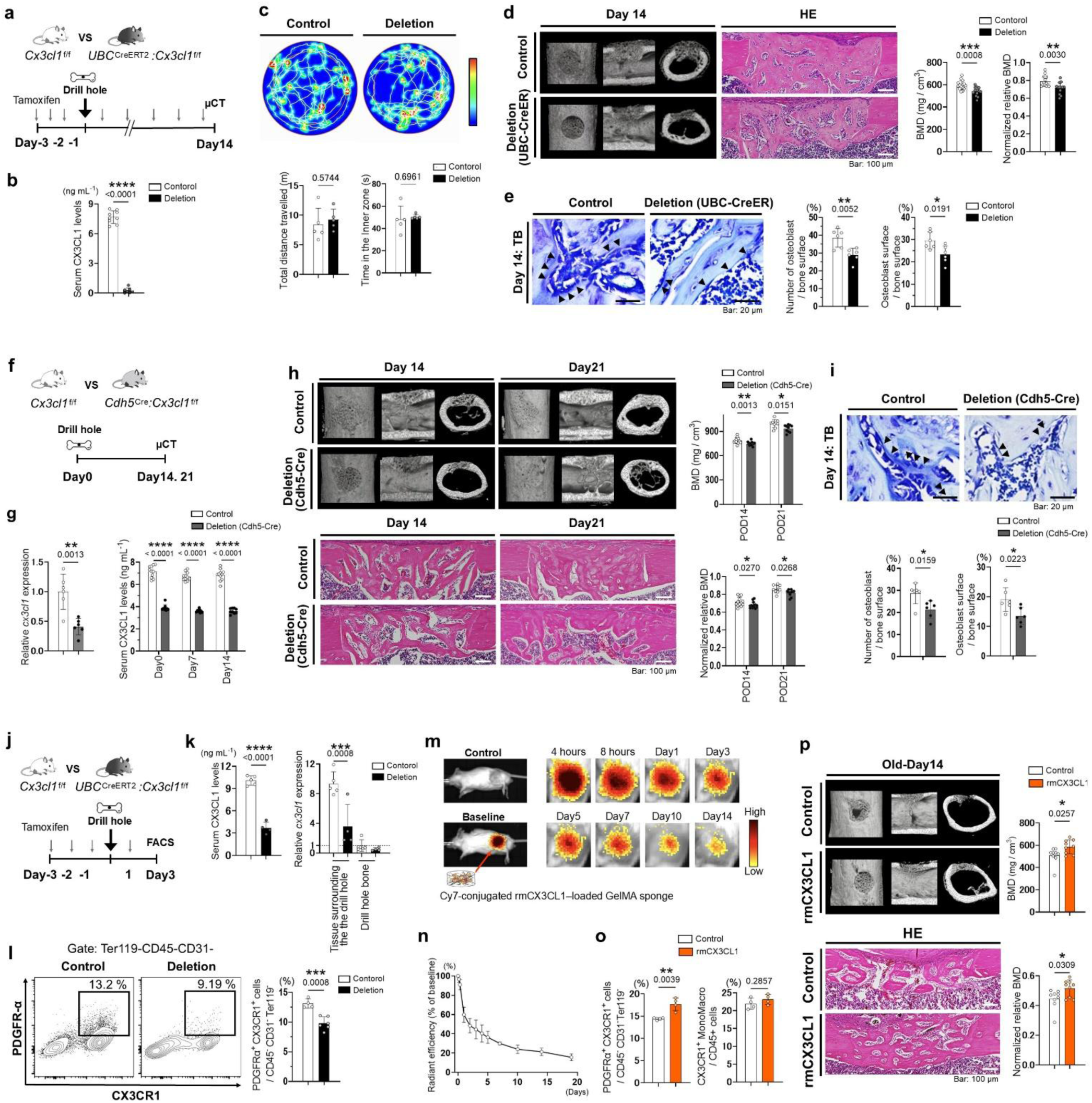
CX3CL1 loss delays bone repair and reduces CX3CR1⁺ osteoprogenitor accumulation a,b, Tamoxifen induced global *Cx3cl1* deletion in UBC-CreERT2; *Cx3cl1* flox/flox mice. a, Timeline. b, Serum CX3CL1 at day 14 after injury. n = 7–9 per group. **c,** Open field analysis of uninjured control and UBC-CreERT2; Cx3cl1 flox/flox mice on day 10. n = 5 per group. **d,e,** Bone repair after global *Cx3cl1* deletion. d, μCT and H&E images with BMD and normalized relative BMD at day 14. Normalized relative BMD = repair region BMD/contralateral cortical BMD. n = 13–15 per group, 5–9 per sex. e, Toluidine blue images and osteoblast histomorphometry. Arrowheads indicate osteoblasts. n = 6 per group. **f,g,** Endothelial *Cx3cl1* deletion in Cdh5-Cre; *Cx3cl1* flox/flox mice. f, Model schematic. g, Local *Cx3cl1* mRNA and serum CX3CL1 at day 14. n = 6 per group for qPCR and n = 9–11 for serum. **h,i,** Bone repair after endothelial *Cx3cl1* deletion. h, μCT and H&E images with BMD and normalized relative BMD at days 14 and 21. n = 13–15 per group at day 14, 5–9 per sex, and n = 9–10 at day 21, 4–6 per sex. i, Toluidine blue images and osteoblast histomorphometry at day 14. n = 6 per group. **j–l,** Global *Cx3cl1* deletion and analysis of CX3CR1⁺PDGFRα⁺ stromal cells at day 3. j, Model schematic. k, Serum CX3CL1 and local *Cx3cl1* mRNA. n = 4–5 per group. l, Flow cytometry analysis of CX3CR1⁺PDGFRα⁺ cells among Ter119⁻CD45⁻CD31⁻ nonhematopoietic cells in adjacent bone tissue. n = 4–6 per group. **m,n,** Longitudinal IVIS imaging of GelMA sponges loaded with Cy7 conjugated rmCX3CL1. Signals were normalized to baseline. n = 4 mice. **o**, Lin⁻PDGFRα⁺CX3CR1⁺ cells and CD45⁺CX3CR1⁺ monocytes/macrophages in injured bone at day 3 after rmCX3CL1 treatment. Lin⁻ cells were Ter119⁻CD45⁻CD31⁻. n = 4 per group. **p,** μCT and H&E analysis with BMD and normalized relative BMD in older male mice after rmCX3CL1 treatment at day 14. n = 9 per group, 23–25 months old. Data are mean ± SD. Student’s t test was used.

Open field testing revealed no significant differences between groups across multiple locomotion parameters, consistent with previous reports using CX3CL1 deficient mice (Fig. 5c and Supplementary Fig. 10a,b) ^39^. μCT analysis at day 14 showed significantly reduced bone mineral density in mice lacking CX3CL1 compared with controls (Fig. 5d). We found a reduction in osteoblast related parameters using bone histomorphometric analysis, whereas osteoclast related parameters were not substantially altered, suggesting that the impaired repair was more closely associated with reduced osteoblast related activity (Fig. 5e and Supplementary Fig. 11a).

As a second approach to assess the effects of CX3CL1 during bone repair, we employed *Cx3cl1* flox/flox mice and induced local deletion using adenovirus mediated Cre recombination at the injury site^40–43^. Deletion efficiency was confirmed by a marked reduction of *Cx3cl1* expression in muscle adjacent to the injury site with no significant change in serum CX3CL1 levels (Supplementary Fig. 12a-e). μCT analysis at days 14 and 21 revealed impaired bone healing in the deletion group compared with controls (Supplementary Fig. 12f,g). Consistent with this, bone histomorphometric analysis demonstrated decreases in osteoblast related parameters, whereas osteoclast related parameters were not significantly altered (Supplementary Fig. 12h,i).

To assess the contribution of endothelial derived CX3CL1 during bone repair, we examined *Cdh5*-Cre mediated deletion of *Cx3cl1*. Deletion efficiency was confirmed by reduced serum CX3CL1 levels together with reduced *Cx3cl1* expression in muscle surrounding the drill hole (Fig. 5f,g). μCT analysis at days 14 and 21 revealed impaired bone healing in the deletion group compared with controls (Fig. 5h). Bone histomorphometric analysis further showed decreases in osteoblast related parameters, whereas osteoclast related parameters were not significantly altered (Fig. 5i and Supplementary Fig. 11b).

We next examined whether CX3CL1 loss affects the accumulation of PDGFRα⁺CX3CR1⁺ osteoprogenitors during repair. Flow cytometric analysis of bone from mice with inducible global deletion of *Cx3cl1* revealed a significant reduction in PDGFRα⁺CX3CR1⁺ osteoprogenitor cells at the injury site in *Cx3cl1* deleted mice (Fig. 5j–l).

Given this reduction, we asked whether local CX3CL1 delivery could enhance the accumulation of CX3CR1^+^ cells in older mice, in which fewer numbers of these cells accumulate after a bone injury (Fig. 3h). Recombinant CX3CL1 was loaded into a gelatin methacrylate-based sponge and implanted at the injury site (Supplementary Fig. 13). In vivo imaging using Cy7 conjugated CX3CL1 confirmed sustained local retention over the early repair period (Fig. 5m,n). Local delivery of recombinant CX3CL1 significantly increased the number of PDGFRα⁺CX3CR1⁺ osteoprogenitors at the injury site compared with controls, whereas CX3CR1⁺ monocyte and macrophage populations were not significantly changed (Fig. 5o). To assess therapeutic potential in older animals, CX3CL1 loaded scaffolds were implanted immediately after injury and bone repair was evaluated by μCT. At day 14, local CX3CL1 delivery improved bone repair in older mice compared with control scaffolds, indicating that CX3CL1 can partially restore impaired bone repair in older mice (Fig. 5p). Together, these findings indicate that CX3CL1 promotes bone repair, at least in part, by supporting the accumulation of PDGFRα⁺CX3CR1⁺ osteoprogenitors at the injury site.

## Discussion

The molecular signals through which muscle regulates bone healing have remained poorly defined^1–3,29^. Here, we show that bone injury induces a localized CX3CL1 response in the muscle vasculature adjacent to the injury site. CX3CL1 expression is enriched in endothelial subsets associated with regenerative activity. These findings indicate that, beyond its previously recognized roles as a structural soft tissue component and a potential source of osteogenic cells, injury adjacent muscle functions as an active signaling niche that regulates osteoprogenitor behavior during repair. Notably, this response was attenuated with aging, suggesting that impaired local muscle to bone communication contributes to age related defects in bone regeneration.

While CX3CL1–CX3CR1 signaling has been studied largely in the context of immune surveillance, neuroimmune interactions, and osteoclast related biology, our findings show that this pathway also engages a nonhematopoietic stromal compartment involved in bone regeneration^17,24–26,44^. CX3CR1 was detected in PDGFRα+ osteoprogenitors, enriched near sites of new bone formation, and functionally linked to proliferative and osteogenic responses to CX3CL1. Thus, in the context of bone repair, CX3CL1 acts locally as a pro regenerative factor. A similar link has been reported in vascular calcification, where CX3CL1–CX3CR1 signaling has been linked to osteogenic transformation of CX3CR1⁺ vascular smooth muscle cells^45^. Although this occurs in a distinct pathological setting, it supports the possibility that this pathway may influence osteogenic differentiation in mesenchymal lineage cells.

An important aspect of our findings is that the CX3CL1 response was highly localized. Endothelial cells represented a substantial fraction of the local CX3CL1⁺ population, consistent with prior studies identifying endothelial cells as a well-recognized source of CX3CL1 under inflammatory or tissue stress conditions^30,31^. Moreover, CX3CL1⁺ cells were preferentially enriched in CD200⁺ endothelial subsets that also expressed CD157 and/or EMCN, markers linked to endothelial regenerative potential and angiogenic osteogenic coupling. This pattern suggests that CX3CL1 may be associated with a repair related endothelial state in injury adjacent muscle^33–35^.

Although CX3CL1–CX3CR1 signaling has been linked to osteoclast lineage cells, our findings suggest that this axis is more closely associated with osteoblast related repair processes in this injury model. The limited osteoclast related changes in our model may also reflect its distinct context, as previous studies examined homeostatic, osteoporotic, or arthritis related bone resorption^24–26,46^.

Injury induced CX3CL1 upregulation in adjacent muscle was attenuated in older mice, accompanied by reduced accumulation of PDGFRα⁺CX3CR1⁺ osteoprogenitors. Local delivery of CX3CL1 increased the number of these osteoprogenitors and partially improved bone repair in older mice, indicating that reduced local CX3CL1 signaling contributes to impaired healing with aging. Together, these findings suggest that impaired muscle to bone signaling is one feature of age-related impairment in bone regeneration.

Communication between neighboring tissues is a fundamental but still incompletely understood feature of regeneration. Our findings identify a CX3CL1 mediated muscle to bone axis that supports bone repair. Defining the roles and mechanisms of adjacent tissue communication will be important for the development of new therapeutic strategies to improve bone healing, particularly when the local regenerative environment is impaired.

## Methods

### Mice

All animal procedures were approved by the Institutional Animal Care and Use Committee of Duke University. *Cx3cr1^eGFP^* (B6.129P-*Cx3cr1^tm1Litt^*/J), UBC-CreERT2 (B6.Cg-*Ndor1^Tg(UBC-cre/ERT2)1Ejb^*/1J), *Pdgfrα*-CreERT2 (B6.129S-*Pdgfra^tm1.1(cre/ERT2)Blh^*/J), *R26^tdTomato^* (B6.Cg-Gt(*ROSA*)*26Sor^tm^*^14^*^(CAG-tdTomato)Hze^*/*J*), *Cdh5*-Cre (B6.FVB-Tg(*Cdh5*-cre)7Mlia/J), and C57BL/6J mice were purchased from Jackson Laboratory. Older C57BL/6J mice were provided by the National Institute on Aging (NIA). Unless otherwise stated, young mice were 3 to 4 months old, and older mice were 23 to 25 months old. *Col1a1*-Cre mice, which express Cre recombinase under the control of the 2.3 kb *Col1a1* promoter in osteoblasts, were obtained from the Mutant Mouse Resource and Research Center at the University of California, Davis^47,48^. *Cx3cl1*-flox mice (C57BL/6JGpt-*Cx3cl1*^em1Cflox^/Gpt) were purchased from GemPharmatech. All mice were bred under specific-pathogen-free conditions. UBC-CreERT2 (B6.Cg-*Ndor1^Tg(UBC-cre/ERT2)1Ejb^*/1J) and *Cdh5*-Cre (B6.FVB-Tg(*Cdh5*-cre)7Mlia/J) were mated with *Cx3cl1*-flox (C57BL/6JGpt-*Cx3cl1*^em1Cflox^/Gpt) to generate UBC-CreERT2; *Cx3cl1*-flox or *Cdh5*-Cre; *Cx3cl1*-flox mice, respectively. *Col1a1*-Cre mice were mated with R26tdTomato mice to generate *Col1a1*-Cre; R26tdTomato mice. *Pdgfrα*-CreERT2 (B6.129S-*Pdgfra^tm1.1(cre/ERT2)Blh^*/J), *Col1a1*-Cre (FVB-Tg(*Col1a1*-cre)1Kry/Mmucd), *R26^tdTomato^* (B6.Cg-Gt(*ROSA*)*26Sor^tm14(CAG-tdTomato)Hze^*/*J*), and *Cx3cr1^eGFP^* (B6.129P-*Cx3cr1^tm1Litt^*/J) mice were mated to generate *Pdgfrα*-CreERT2; *R26^tdTomato^*; *Cx3cr1^eGFP^* or *Col1a1*-Cre; *R26^tdTomato^*; *Cx3cr1^eGFP^* mice, respectively. Genotyping PCR for each mouse strain was performed based on protocols provided by the respective repositories or suppliers, with minor modifications. To induce Cre-mediated recombination, tamoxifen (Sigma-Aldrich) was dissolved in corn oil (Sigma-Aldrich) and administered intraperitoneally at a dose of 80 μg g⁻¹ body weight. UBC-CreERT2 mice received tamoxifen on days −3, −2, and −1 before injury and every other day after injury, whereas *Pdgfra*-CreERT2 lineage tracing mice received tamoxifen on days −10, −9, and −8 before injury.

### Drill hole surgery

Mice underwent drill hole surgery under anesthesia. For the drill hole injury model, the left femur was operated on, while the right femur served as an internal control^49,50^. A small incision was made in the lateral skin at the mid femur. After blunt separation of the surrounding muscle, the anterior surface of the mid femur was exposed. A drill hole injury was created in the mid femur using a micro drill equipped with a 0.8 mm dental carbide bur (Komet Dental, Germany). The muscle and skin were subsequently closed with nylon sutures. Mice were euthanized at the indicated time points after surgery for tissue collection.

### Local adenoviral Cre mediated deletion of *Cx3cl1*

Previous studies from our group and others have shown that local adenoviral Cre delivery effectively induces recombination at skeletal injury sites^40–43^. Local deletion of *Cx3cl1* was therefore induced by adenoviral delivery of Cre recombinase to the bone injury site. Briefly, *Cx3cl1* flox/flox mice received a local injection of adenovirus expressing Cre recombinase (Vector Biolabs) into the femoral drill hole site immediately after injury. Control mice received an equivalent dose of adenovirus expressing GFP, referred to as Ad-control. Tissue surrounding the drill hole, including adjacent muscle and repair tissue, was collected, and local recombination and *Cx3cl1* deletion were assessed by qPCR, RNAscope in situ hybridization, and immunofluorescence staining.

### Local delivery of rmCX3CL1

For local CX3CL1 treatment, recombinant mouse CX3CL1, rmCX3CL1 (R&D Systems, Cat. 472-FF), was delivered using gelatin methacryloyl (GelMA) sponges (Supplementary Fig. 13). GelMA was synthesized by methacrylation of gelatin. GelMA hydrogels were prepared by photopolymerization of GelMA in the presence of the photoinitiator LAP under UV light. The resulting hydrogels were frozen at −80 °C and freeze dried to generate porous GelMA sponges. GelMA synthesis was characterized by FTIR and ¹H NMR, and the porous architecture of the freeze dried GelMA sponge was assessed by scanning electron microscopy. GelMA sponges were loaded with rmCX3CL1 at a dose of 20 ng per mouse and implanted into the femoral drill hole site immediately after injury. Control mice received PBS loaded GelMA sponges. For retention analysis, GelMA sponges were loaded with Cy7 labeled rmCX3CL1 and implanted into the femoral drill hole site.

### Three-dimensional micro-computed tomography analysis

Femurs were analyzed by micro computed tomography using the ScanXmate L090H system (Comscantecno, Yokohama, Japan). Three-dimensional reconstructions were generated using the TRI 3D Bon FSC system (RATOC System Engineering, Santa Clara, CA, USA), as previously described^49–51^. A cylindrical region of interest was positioned within the cortical bone at the injury site and spanned the cortical thickness from the endosteal to periosteal surface. Regenerated bone was evaluated by BMD and normalized BMD, with normalization performed relative to the contralateral cortical bone.

### Real time PCR analysis

Total RNA was isolated from the indicated tissues, including adjacent muscle, repair tissue, and organs, and reverse transcribed into cDNA. Quantitative PCR was performed using TaqMan Gene Expression Assays, Thermo Fisher Scientific, according to the manufacturer’s instructions. The following assays were used: mouse *Cx3cl1*, Mm00436454_m1, and mouse *Gapdh*, Mm99999915_g1. *Gapdh* was used as the endogenous control, and relative *Cx3cl1* expression was calculated after normalization to *Gapdh*.

### Histological analysis

Paraffin embedded femurs were fixed in 4% paraformaldehyde for 24–48 h at 4 °C and subsequently decalcified in 10% ethylenediaminetetraacetic acid for approximately 14 days. Decalcified tissues were dehydrated through graded ethanol concentrations and embedded in paraffin. Serial sections (7 μm) of the femurs were prepared using a microtome (Leica Microsystems, RM2235) and subjected to hematoxylin and eosin, TRAP, and toluidine blue staining. Bone histomorphometric analyses were performed using ImageJ software (National Institutes of Health) in accordance with standardized guidelines of the American Society for Bone and Mineral Research, as previously described^52^. For cryosectioning, femurs were fixed in 4% PFA (paraformaldehyde) in PBS at 4 °C overnight and subsequently incubated overnight in a 15% to 30% sucrose gradient in PBS. Tissues were embedded in Tissue Tek O.C.T. compound (Sakura Finetek). Cryosections were prepared at a thickness of 14 μm using a CryoJane Tape Transfer System and a CM1950 cryostat (Leica Microsystems). For immunostaining, sections were incubated overnight at 4 °C with the following primary antibodies: rabbit anti CX3CL1 polyclonal antibody (1:300, Abcam, Cat. ab25088), rat anti CD31 monoclonal antibody (1:300, clone MEC 13.3, BD Biosciences, Cat. 550274), rat anti CD45 monoclonal antibody (1:1000, clone 30 F11, Cell Signaling Technology, Cat. 55307), rat anti mouse F4/80 monoclonal antibody (1:200, clone A3 1, Bio Rad, Cat. MCA497R), goat anti mouse ALP polyclonal antibody (1:100, R&D Systems, Cat. AF2910). Sections were subsequently incubated with species appropriate secondary antibodies for 3 h at room temperature, including Alexa Fluor 647 conjugated goat anti rat IgG (1:500, Abcam, Cat. ab150167), Alexa Fluor 594 conjugated goat anti rat IgG (1:500, Abcam, Cat. ab150160), and APC conjugated goat anti rabbit IgG (1:500, Thermo Fisher Scientific, Cat. A10931). Nuclei were counterstained with DAPI (Abcam, Cat. ab104139).

### RNA Scope

Tissues for RNAscope were fixed in 4% paraformaldehyde, decalcified in Morse’s solution for optimal RNA preservation, dehydrated, embedded in paraffin, and sectioned at 7 μm thickness for RNAscope analysis, as previously described^53,54^. RNAscope in situ hybridization was performed using the RNAscope Multiplex Fluorescent V2 Assay (Advanced Cell Diagnostics, Cat. 323100). Probes targeting Cx3cl1, Advanced Cell Diagnostics, Cat. 426211 C1, and Cx3cr1, Advanced Cell Diagnostics, Cat. 314221 C2, were used to visualize their spatial expression patterns. Signal amplification was performed according to the manufacturer’s protocol. Fluorescent signals were developed using Vivid 570 (Tocris, Cat. 7526), and Vivid 650 (Tocris, Cat. 7527), for 30 min at room temperature. Nuclei were counterstained with DAPI (Abcam, Cat. ab104139).

### Image processing and quantifications

Immunostained samples were imaged with a Zeiss LSM880 (Carl Zeiss). The images were processed using Zen blue (Carl Zeiss), Adobe Illustrator (Adobe), ImageJ2 (v.1.53u; Fiji; http://fiji.sc/). Most figures show maximum-intensity projections unless otherwise indicated. Tiles scan overview images of bone were taken with 20x lens. Brightness-contrast modifications were uniformly applied to the original images to improve image quality and visualization. Marker positive cells were quantified using marker signals and corresponding tissue regions, as specified in the corresponding figures and figure legends. Images shown are representative and were obtained from multiple independent experiments, each performed using identical laser excitation and confocal detector settings within the same experiment. At least three mice were analyzed per condition.

### FACS analysis of injured bone and peri-hole tissue

Soft tissues were carefully removed from femurs (Fig. 1a and Supplementary Fig. 1c). Femurs were mechanically triturated and incubated with 0.2% collagenase type2 and 50 μg/mL DNase I in αMEM for 30 min at 37 °C with gentle agitation, followed by filtration through a 70-μm cell strainer on ice to generate single-cell suspensions.

For digestion of tissue surrounding the drill hole, tissues were carefully dissected and minced into small fragments on ice. Samples were transferred to gentleMACS C tubes containing digestion buffer (αMEM supplemented with collagenase II [800 U/mL] and dispase [2.4 U/mL]) and enzymatically dissociated using a gentleMACS Dissociator (Miltenyi Biotec) for approximately 60 min at 37 °C with gentle agitation. Following enzymatic digestion, cell suspensions were filtered through a 70-μm cell strainer and further processed for debris removal using a commercial debris removal solution according to the manufacturer’s instructions (Miltenyi Biotec, Cat# 130-096-334).

Single-cell suspensions were then subjected to red blood cell lysis, followed by passage through a 40-μm mesh filter and washing with FACS buffer consisting of DPBS supplemented with 2.0% heat-inactivated fetal bovine serum and 2.5 mM EDTA.

Cells were incubated with Fc receptor blocking reagent (anti-mouse CD16/32; Invitrogen) for 10 min on ice and subsequently stained with fluorophore-conjugated antibodies against mouse CD45 (30-F11), CD11b (M1/70), F4/80 (BM8), Ly6G (1A8), CD3e (145-2C11), CD4 (GK1.5), CD8a (53-6.7), CD11c (N418), NK1.1 (PK136), CD45R/B220 (RA3-6B2), CX3CR1 (SA011F11), PDGFRα (APA5), CD31 (390), Ter119 (TER-119), CD200 (OX-90), CD157 (BP-3), Endomucin (V.7C7), and CX3CL1 (126315). Antibodies were purchased from BioLegend, R&D Systems, Invitrogen, or BD Biosciences. Dead cells were excluded using the LIVE/DEAD Fixable Near-IR Dead Cell Stain Kit, Thermo Fisher Scientific. Details of antibodies and viability dyes are provided in Supplementary Table 1. Cells were washed and resuspended in FACS buffer for analysis. Flow cytometric analyses were performed on a Cytek Aurora (Cytek Biosciences). Cell sorting was conducted using a Beckman Coulter Astrios, and post-sort purity was confirmed. Data were analyzed using FlowJo software (BD Biosciences, v10.10.0).

### Cell proliferation and viability assays

Femurs were mechanically dissociated and incubated with 0.2% collagenase type2 and 50 μg/mL DNase I in αMEM for 30 min at 37 °C with gentle agitation. Digested tissues were filtered through a 70 μm cell strainer on ice to generate single cell suspensions. CD45-depleted bone-derived mesenchymal stromal cells were obtained by MACS-based CD45 depletion and expanded under hypoxic culture conditions as previously described^38^. Passage 3 cells were used for EdU incorporation and CellTiter-Glo assays. For the EdU assay, cells were seeded on chambered glass slides at approximately 3.5 × 10⁴ cells/cm². After 24 h, cells were treated with PBS vehicle control or recombinant mouse CX3CL1 (rmCX3CL1, 100 ng/ml). Twenty-four hours after treatment, EdU was added to the culture medium, and cells were incubated for 4 h. Cells were then fixed and stained using an EdU Click reaction according to the manufacturer’s protocol, with DAPI used for nuclear counterstaining. For quantification, nine 20× fields of view were acquired from each independent sample, and the average value was used as one biological replicate for statistical analysis. For the CellTiter-Glo assay, passage 3 CD45⁻ mesenchymal stromal cells were seeded in 96 well plates at 1.0 × 10⁴ cells per well. After 24 h, cells were treated with PBS vehicle control or rmCX3CL1 (100 ng/mL). Cell viability and proliferation were assessed at 0, 24 and 48 h after treatment using the CellTiter-Glo luminescent cell viability assay according to the manufacturer’s protocol. ATP dependent luminescence signals were measured using a plate reader.

### Osteogenic differentiation assays

CD45⁻ mesenchymal stromal cells isolated from bone tissue were expanded under hypoxic conditions, and passages 3 to 5 cells were used for osteogenic differentiation assays^38^. Cells were seeded in culture plates at approximately 1.5 × 10⁴ to 3.0 × 10⁴ cells/cm². On day 1 after seeding, cells were cultured in osteogenic medium with PBS vehicle control or rmCX3CL1 (100 ng/mL). In a subset of ALP experiments, cells were cultured in αMEM without osteogenic medium. Medium was changed every 3 days. Cells were fixed on day 7 for ALP staining and on day 14 for Alizarin Red staining. Staining was performed according to standard protocols, and staining intensity was quantified using a plate reader.

### Serum collection and ELISA

Blood was collected at the indicated time points and serum was isolated by centrifugation. Serum CX3CL1 concentrations were measured using a mouse CX3CL1 ELISA kit (R&D Systems, Cat. MCX310), according to the manufacturer’s protocol. Concentrations were calculated from a standard curve.

### Open field test

Open field testing was performed in uninjured *Cx3cl1* flox/flox and UBC CreERT2; *Cx3cl1* flox/flox mice on day 10 of the tamoxifen treatment timeline using an open field arena equipped with a video tracking system. Mice were placed in the arena and allowed to move freely during the recording period.

Locomotor activity was recorded and analyzed using ANY maze software (Stoelting Co). Total distance traveled and time spent in the inner zone were quantified from the tracking data. Movement traces and heatmaps were generated from the recorded tracking data.

### Statistical analysis

Statistical analyses were performed using GraphPad Prism V11 and StatFlex V7. Data are presented as mean ± SD unless otherwise indicated. For lineage tracing quantification, data are presented as mean ± 95% CI where specified. Comparisons between two groups were performed using two tailed unpaired Student’s t tests. For comparisons among multiple groups, one way ANOVA followed by Tukey’s multiple comparisons test was used. P values < 0.05 were considered statistically significant. Each dot represents an individual mouse or independent biological sample unless otherwise indicated.

## Acknowledgements

We thank Tomoko Towatari and all members of the Department of Orthopedics for their constructive input and discussions. We also thank the Flow Cytometry Core, the Light Microscopy Core Facility, and the Center for Electron Microscopy and Nanoscale Technology at Duke University for their technical expertise and assistance.

## Author Contributions

K.I. and B.A.A. conceived the study. K.I. led the experimental work, analyzed and interpreted the data, and wrote the manuscript. C.M., E.S., P.N., T.N., M.I., J.H., X.M., M.N., N.A., K.A., P.V., and V.P. contributed to experiments, data acquisition, data analysis, and interpretation. T.S., S.V., Y.Y., and B.A.A. provided scientific input, resources, and critical feedback. B.A.A. revised the manuscript and supervised the study. All authors reviewed and approved the manuscript.

---

**Supplementary Fig. 1.**
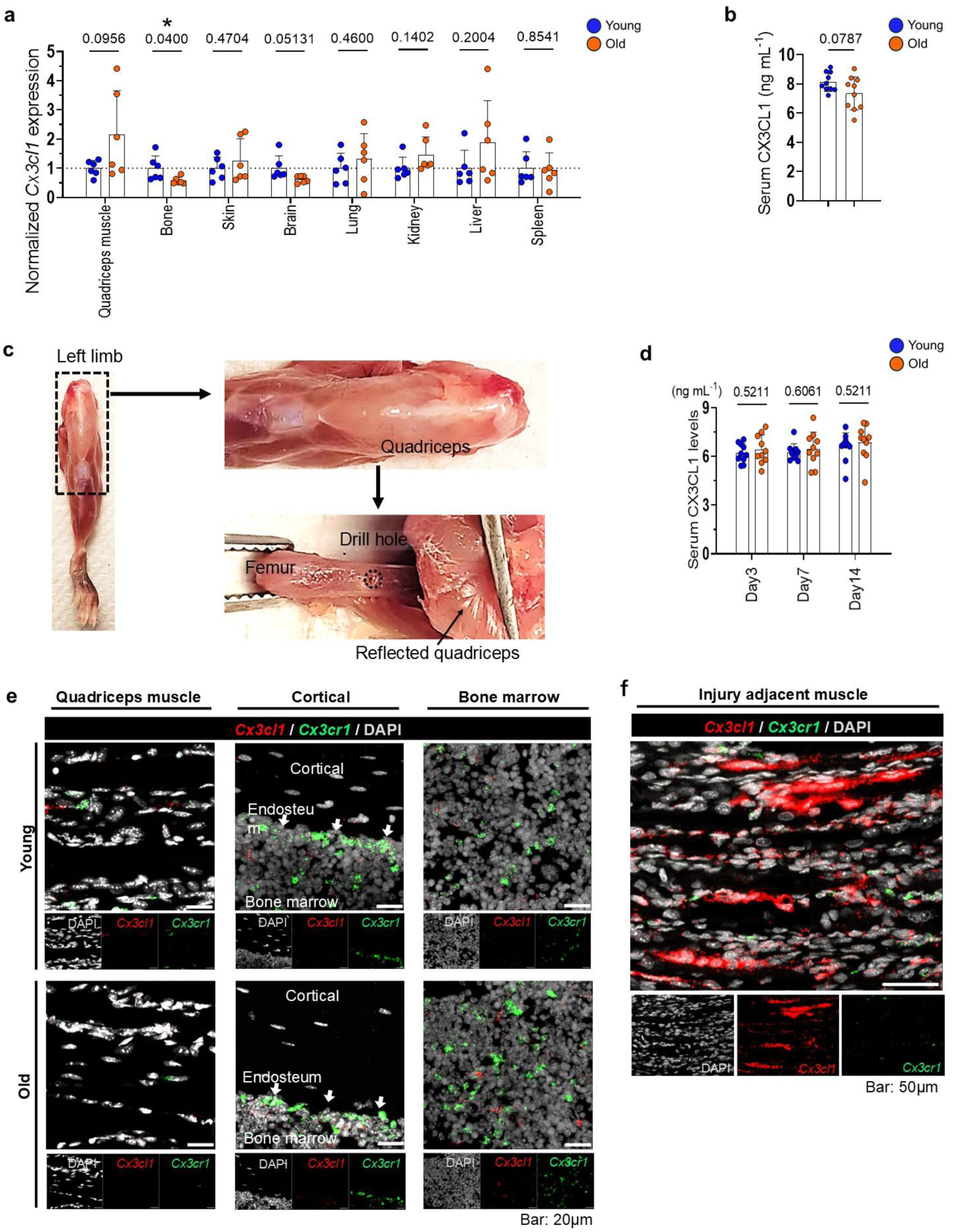
CX3CL1 expression at homeostasis and after injury in young and older mice. a, Relative *Cx3cl1* mRNA expression in the indicated tissues from uninjured young and older mice under homeostatic conditions. Expression was measured by qPCR, normalized to *Gapdh*, and expressed relative to the young group for each tissue. n = 6 mice per age group, including 3 male and 3 female mice. b, Serum CX3CL1 levels in uninjured young and older mice under homeostatic conditions. n = 10 mice per age group, including 5 male and 5 female mice. c, Gross anatomical reference images showing the femoral drill hole and adjacent quadriceps muscle during tissue collection. d, Serum CX3CL1 levels in young and older mice at days 3, 7, and 14 after femoral drill hole injury. n = 10 mice per age group, including 5 male and 5 female mice. e, Representative RNAscope in situ hybridization images of quadriceps muscle adjacent to cortical bone, cortical bone, and bone marrow regions in uninjured young and older mice. *Cx3cl1* is shown in red, *Cx3cr1* in green, and DAPI in gray. Images are representative of n = 5 mice per group. f, Representative RNAscope in situ hybridization images of quadriceps muscle adjacent to the injury site in young mice at day 7 after femoral drill hole injury. Images are representative of n = 5 mice. Young mice were 3 to 4 months old, and older mice were 23 to 25 months old. Data are presented as mean ± SD. Each dot represents an individual mouse or independent sample. Statistical significance was determined using Student’s t test. *P < 0.05.

**Supplementary Fig. 2.**
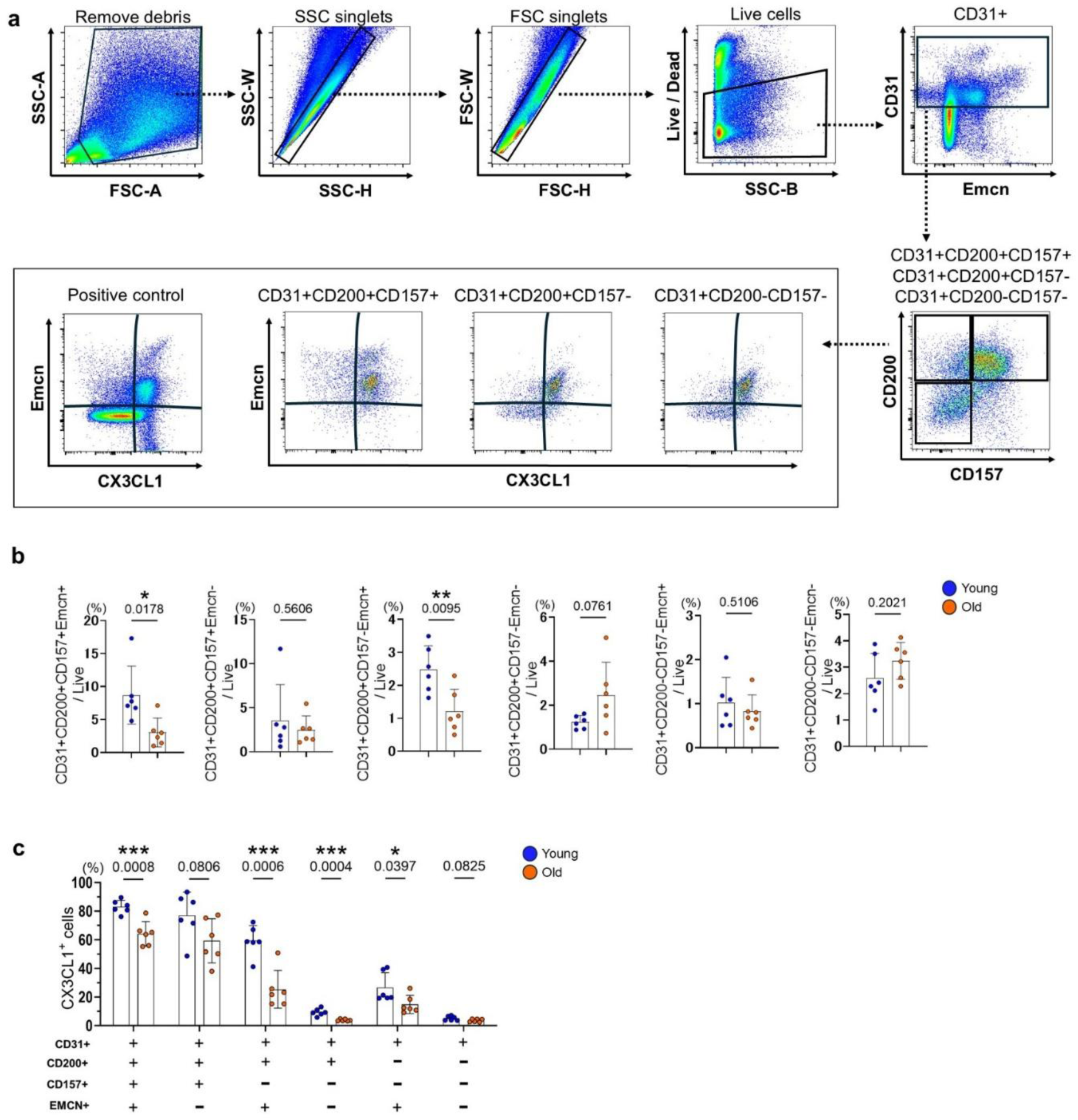
Endothelial subsets in injury adjacent muscle of young and older mice. a, Representative flow cytometry gating strategy for endothelial cell subsets, including CD31⁺CD200⁺CD157⁺EMCN⁺, CD31⁺CD200⁺CD157⁺EMCN⁻, CD31⁺CD200⁺CD157⁻EMCN⁺, CD31⁺CD200⁺CD157⁻EMCN⁻, CD31⁺CD200⁻CD157⁻EMCN⁺ and CD31⁺CD200⁻CD157⁻EMCN⁻ endothelial subsets. b, Quantification of endothelial subsets isolated from muscle tissue surrounding the injured bone in young and older mice at day 5 after injury. Each subset is shown as a percentage of live cells. n = 6 mice per group, including 3 male and 3 female mice. c, Percentage of CX3CL1⁺ cells within each gated endothelial subset in muscle tissue surrounding the injured bone in young and older mice at day 5 after injury. Endothelial subsets were defined by CD200, CD157, and EMCN expression within the live CD31⁺ population. Data are presented as mean ± SD. Statistical significance was determined using Student’s t test.

**Supplementary Fig. 3.**
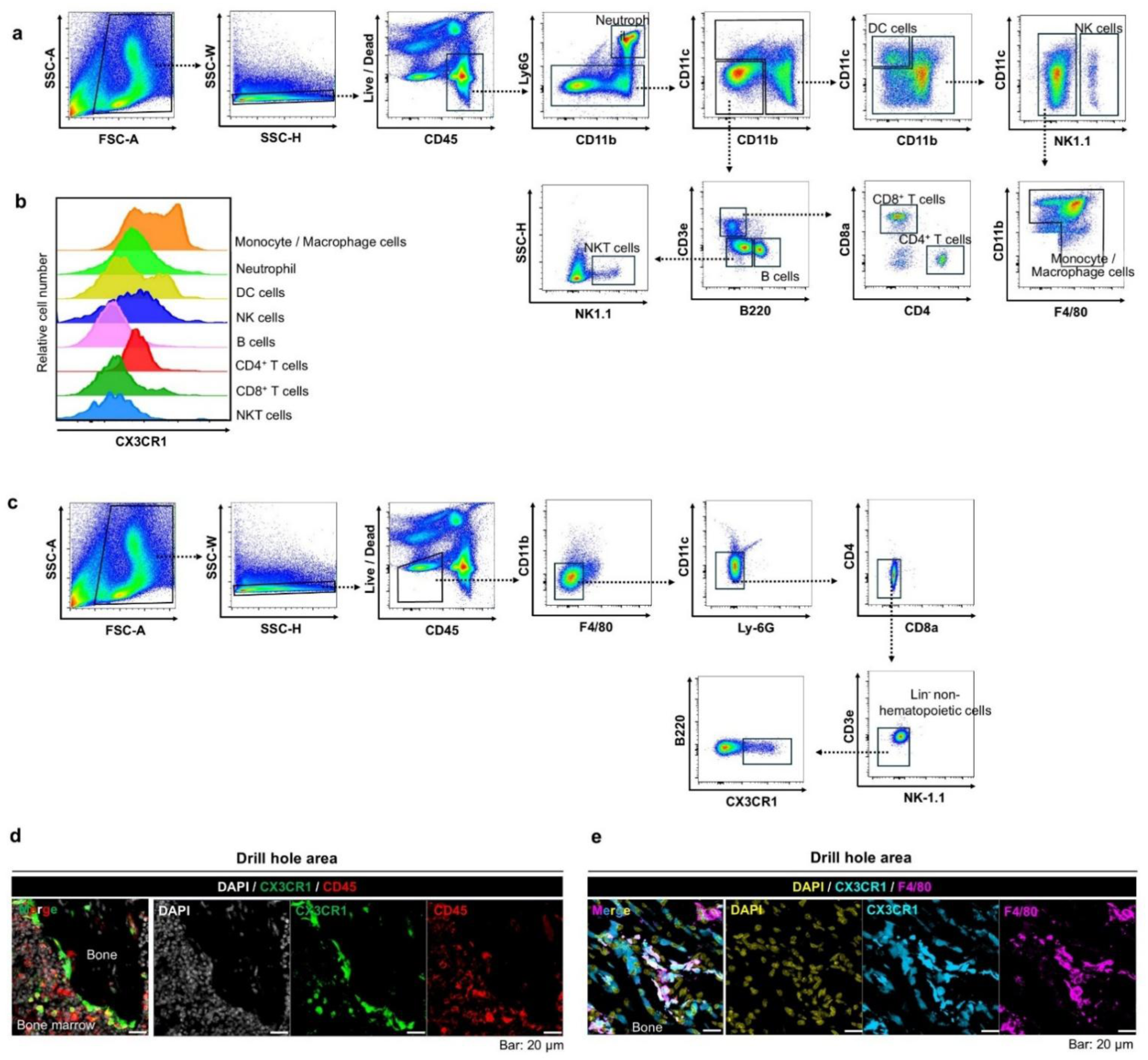
Characterization of CX3CR1 positive hematopoietic and nonhematopoietic cells after bone injury. a–c, Flow cytometry gating strategy for immune cell subpopulations and lineage negative nonhematopoietic cells isolated from bone tissue surrounding the drill hole after femoral drill hole injury. a, Representative flow cytometry plots showing the gating strategy for immune cell subpopulations, including neutrophils, monocyte/macrophage cells, NK cells, DCs, NKT cells, B cells, CD8⁺ T cells, and CD4⁺ T cells. b, Representative overlaid histograms of CX3CR1 expression in each gated population. c, Representative flow cytometry plots showing the gating strategy for Lin⁻CX3CR1⁺ cells. d,e, Representative immunofluorescence images of bone tissue surrounding the drill hole in young *Cx3cr1* GFP/+ reporter mice at day 7 after injury. d, CD45 (red), CX3CR1 GFP reporter signal (green), and DAPI (gray). e, F4/80 (magenta), CX3CR1 GFP reporter signal (cyan), and DAPI (yellow). Images are representative of n = 3 mice per group.

**Supplementary Fig. 4.**
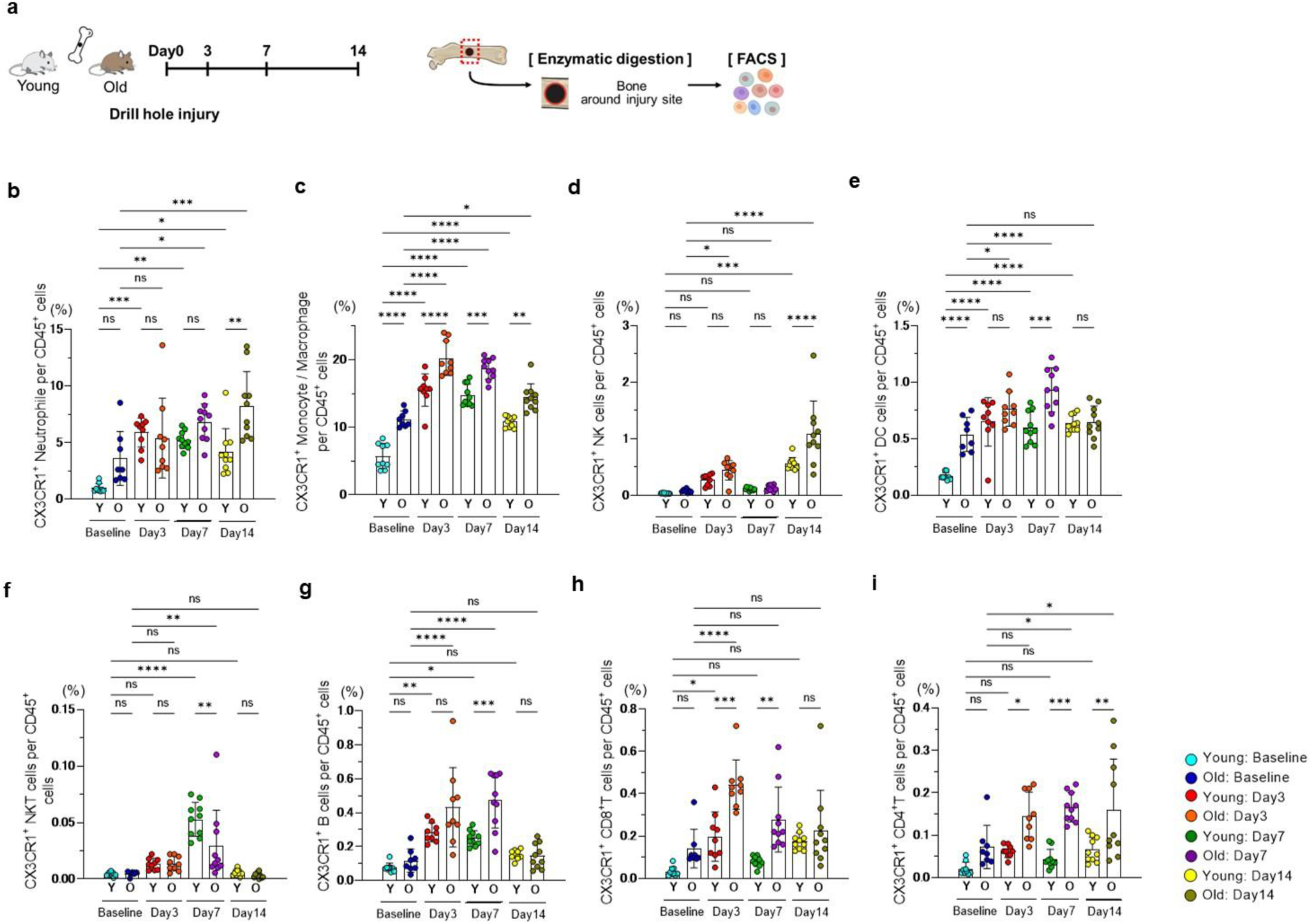
CX3CR1 positive immune cell subpopulations after bone injury in young and older mice. a, Schematic of the experimental design for flow cytometry analysis of CX3CR1⁺ cells in immune cell subpopulations isolated from uninjured femurs or from bone tissue surrounding the drill hole in young and older mice at days 3, 7, and 14 after injury. b–i, Quantification of CX3CR1⁺ cells in each immune cell subpopulation. b, Neutrophils. c, Monocytes/macrophages. d, NK cells. e, DCs. f, NKT cells. g, B cells. h, CD8⁺ T cells. i, CD4⁺ T cells. Quantification shows CX3CR1⁺ cells in each subpopulation as a percentage of total CD45⁺ cells. n = 8–10 mice per group, including equal numbers of male and female mice. Data are mean ± SD. Statistical significance was determined using one-way ANOVA followed by Tukey’s multiple comparisons test. *P < 0.05, **P < 0.01, ***P < 0.001, ****P < 0.0001.

**Supplementary Fig. 5.**
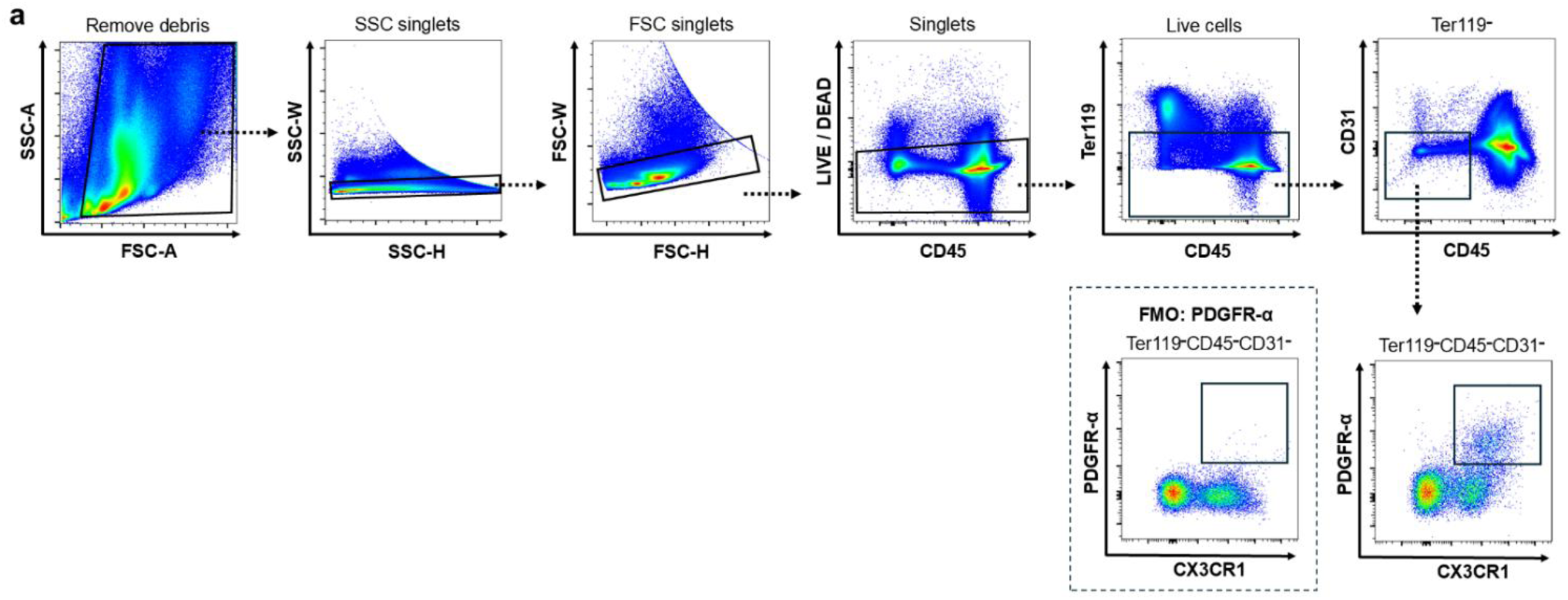
Gating strategy for CX3CR1⁺PDGFRα⁺ stromal cells. a, Representative flow cytometry gating strategy for CX3CR1⁺ cells within PDGFRα⁺ lineage negative nonhematopoietic cells isolated from bone tissue surrounding the drill hole after femoral drill hole injury. Fluorescence minus one (FMO) controls were used to define the PDGFRα⁺ gate.

**Supplementary Fig. 6.**
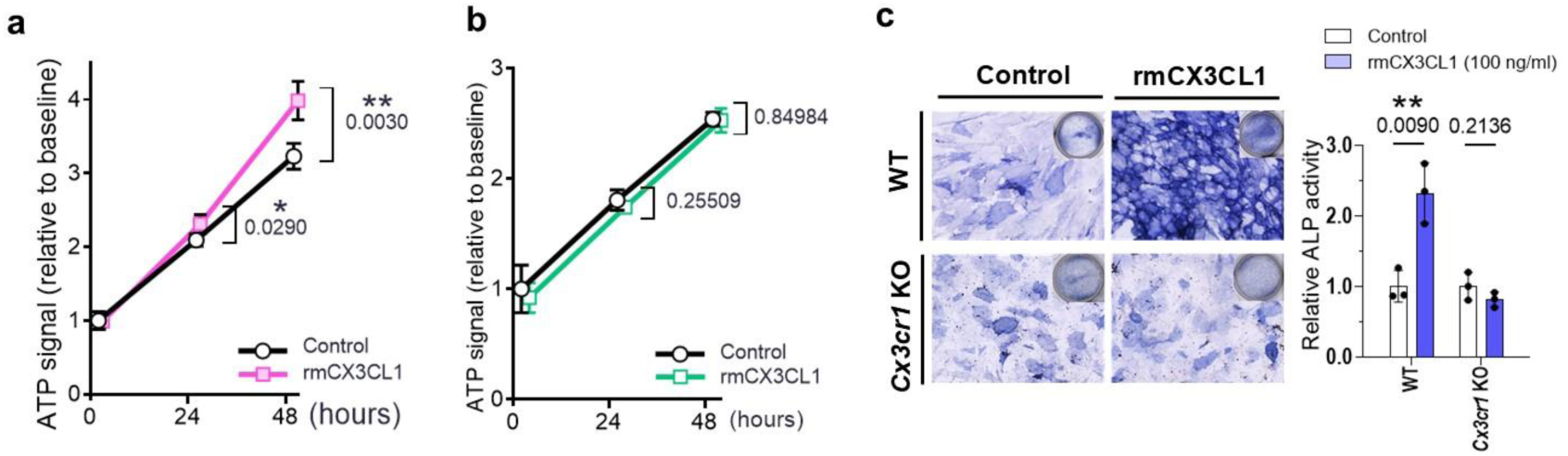
CX3CL1 promotes mesenchymal stromal cell viability through CX3CR1. a, b, Cell viability analysis at 24 and 48 h in CD45 depleted mesenchymal stromal cells isolated from bone tissue of young littermate WT and *Cx3cr1* GFP/GFP mice under control or rmCX3CL1 (100 ng/ml) treated conditions. a, littermate WT cells. b, *Cx3cr1* GFP/GFP cells. c, Representative staining images and quantification of CD45 depleted mesenchymal stromal cells cultured in 20% FBS medium without osteogenic medium under control or rmCX3CL1 treated conditions. n = 3 mice per group. Data are presented as mean ± SD. Each dot represents an individual mouse or independent sample. Statistical significance was determined using Student’s t test. *P < 0.05, **P < 0.01.

**Supplementary Fig. 7.**
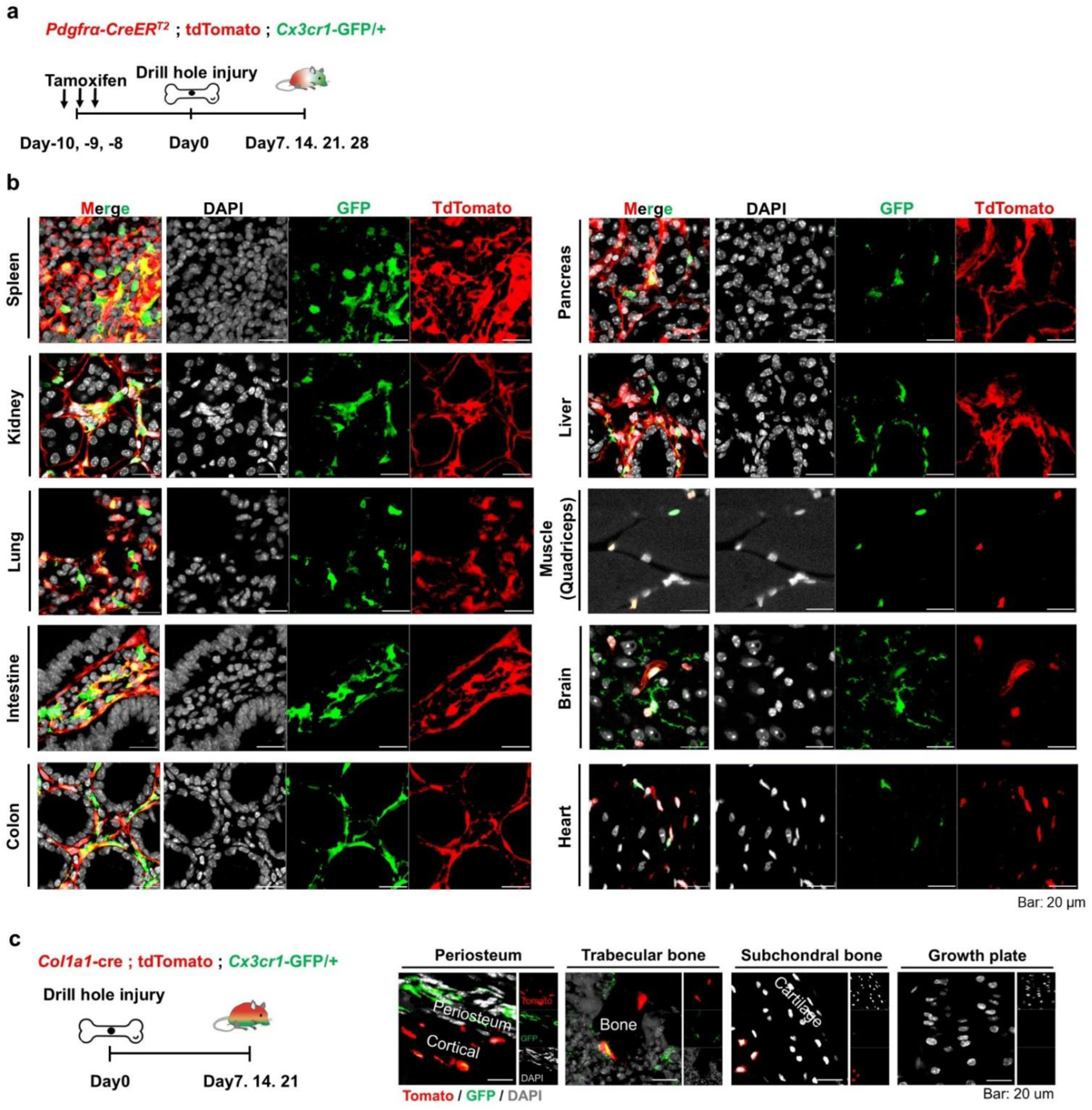
Reporter strategies and tissue distribution of CX3CR1-GFP⁺ cells. a, Experimental timeline for lineage tracing in *Pdgfra*-CreERT2; tdTomato; *Cx3cr1* GFP/+ mice. Tamoxifen was administered before femoral drill hole injury. b, Representative confocal images showing tdTomato and CX3CR1 GFP reporter signals in multiple organs from *Pdgfra-CreERT2*; tdTomato; *Cx3cr1* GFP/+ mice at day 7 after femoral drill hole injury. tdTomato is shown in red, CX3CR1 GFP reporter signal in green, and DAPI in gray. Images are representative of n = 3 mice. c, Experimental timeline and representative fluorescence images of contralateral femoral bone from *Col1a1*-Cre; tdTomato; *Cx3cr1* GFP/+ mice at day 7 after injury. Images are representative of n = 3 mice.

**Supplementary Fig. 8.**
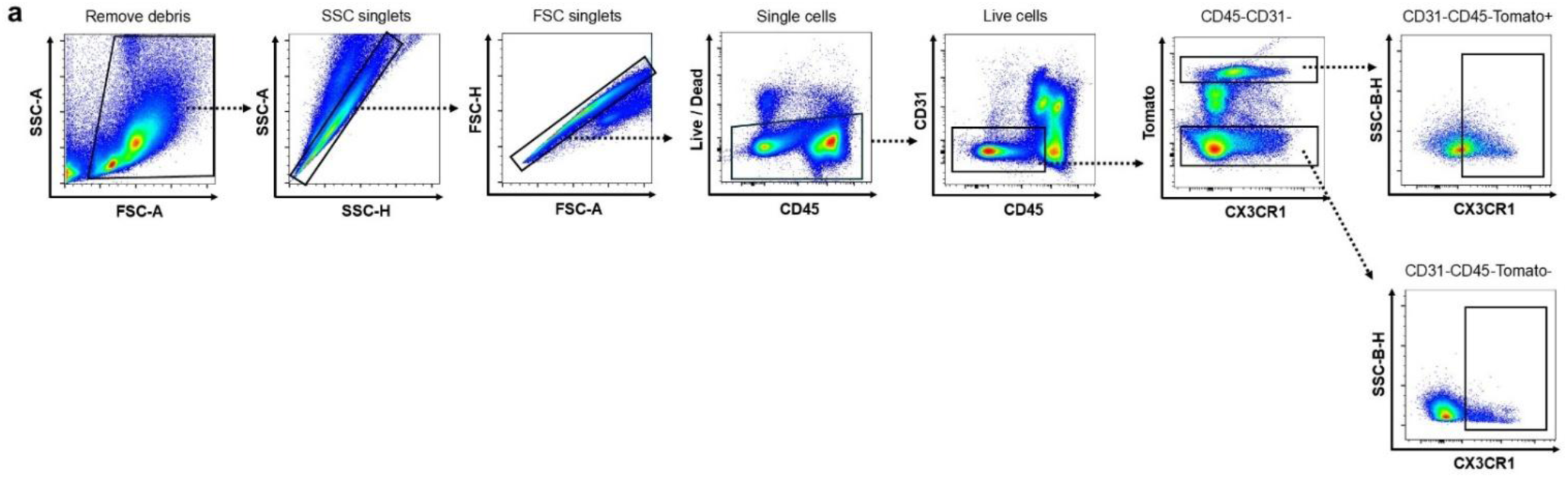
Gating strategy for CX3CR1 positive cells in Col1a1 lineage stromal populations. a, Representative flow cytometry gating strategy for CX3CR1⁺ cells within tdTomato⁺ and tdTomato⁻ CD45⁻CD31⁻ cells isolated from uninjured bone tissue in *Col1a1*-Cre; tdTomato mice. tdTomato⁺ and tdTomato⁻ populations were gated within the CD45⁻CD31⁻ fraction, followed by identification of CX3CR1⁺ cells within each population.

**Supplementary Fig. 9.**
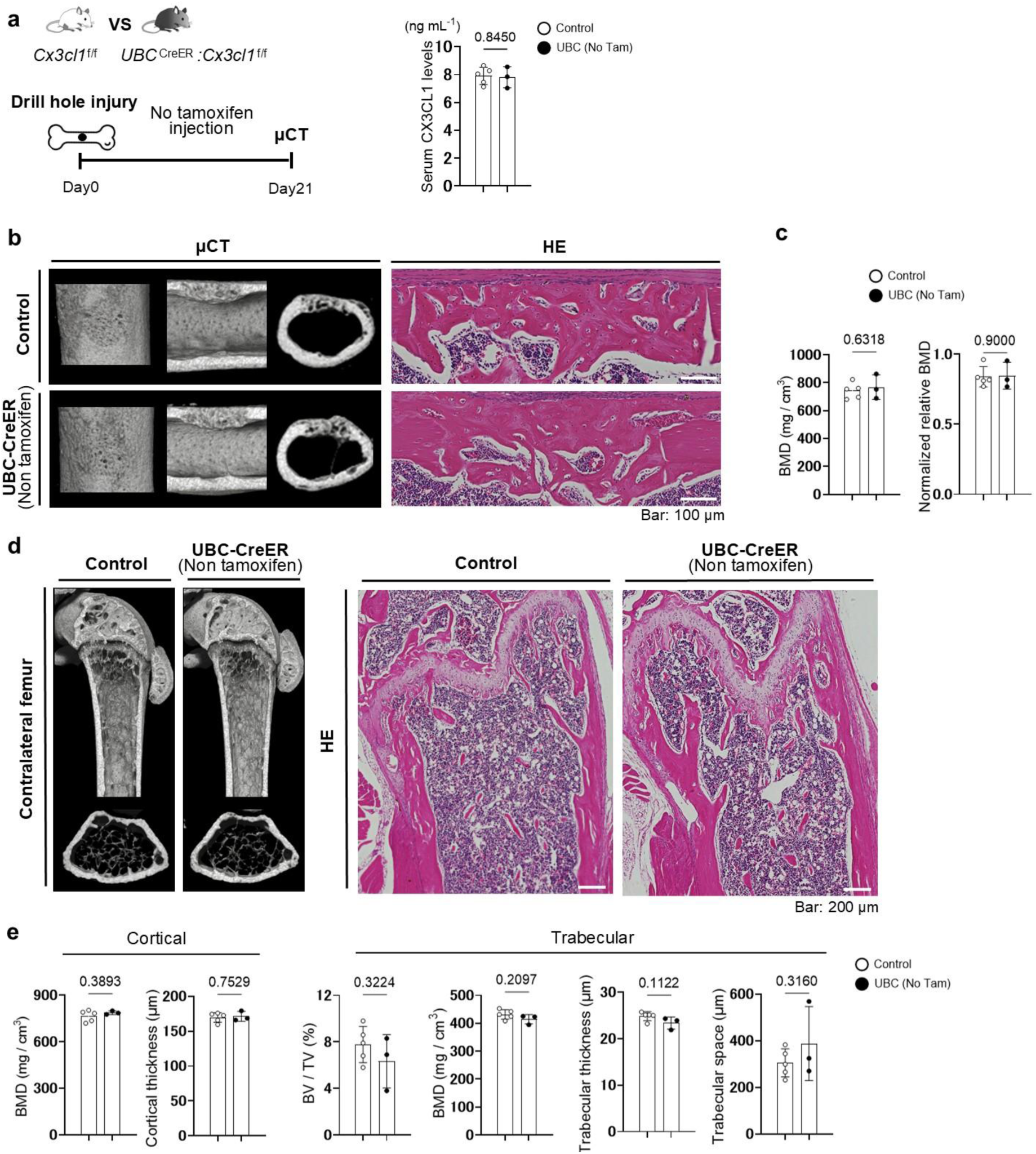
Bone repair without tamoxifen induction in UBC-CreERT2; *Cx3cl1* flox/flox mice. a, Experimental timeline and serum CX3CL1 levels in young UBC-CreERT2; *Cx3cl1* flox/flox mice without tamoxifen induction. Mice underwent femoral drill hole injury without tamoxifen administration and were sacrificed on day 21. Right, serum CX3CL1 levels at day 21. n = 3–5 mice per group. b, Representative μCT images and H&E stained sections of the repair region in UBC-CreERT2; *Cx3cl1* flox/flox mice without tamoxifen induction at day 21 after femoral drill hole injury. c, Quantification of μCT based bone mineral density (BMD) and normalized relative BMD in the repair region at day 21. Normalized BMD was calculated as repair region BMD divided by contralateral cortical BMD. n = 3–5 mice per group. d, Left, Representative μCT reconstructions of the contralateral distal femur in UBC-CreERT2; *Cx3cl1* flox/flox mice without tamoxifen induction at day 21 after femoral drill hole injury. Right, Representative H&E stained sections of the contralateral distal femur at day 21. e, Quantification of cortical and trabecular bone parameters in the contralateral distal femur by microCT at day 21. n = 3–5 mice per group. Data are presented as mean ± SD. Each dot represents an individual mouse or independent sample. Statistical significance was determined using Student’s t test.

**Supplementary Fig. 10.**
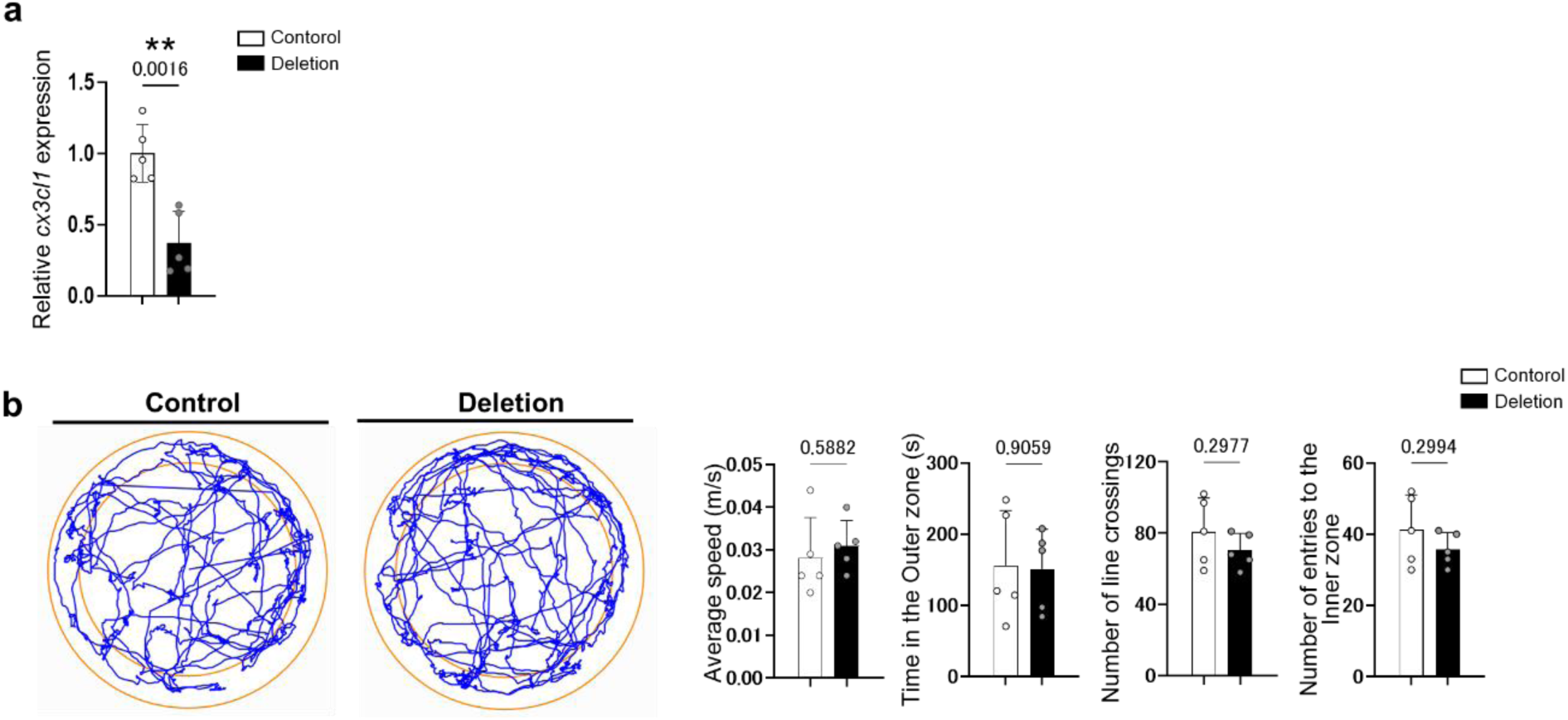
Brain *Cx3cl1* expression and open field behavior after tamoxifen induced global *Cx3cl1* deletion. a, Quantification of brain *Cx3cl1* mRNA expression in young uninjured *Cx3cl1* flox/flox and UBC-CreERT2; *Cx3cl1* flox/flox mice on day 10 of the tamoxifen treatment timeline. *Cx3cl1* expression was assessed by qPCR. n = 5 mice per group. b, Representative open field movement traces showing locomotor activity in uninjured *Cx3cl1* flox/flox and UBC-CreERT2; *Cx3cl1* flox/flox mice on day 10 of the tamoxifen treatment timeline. Open field parameters were quantified as indicated. n = 5 mice per group. Data are presented as mean ± SD. Each dot represents an individual mouse or independent sample. Statistical significance was determined using Student’s t test.

**Supplementary Fig. 11.**
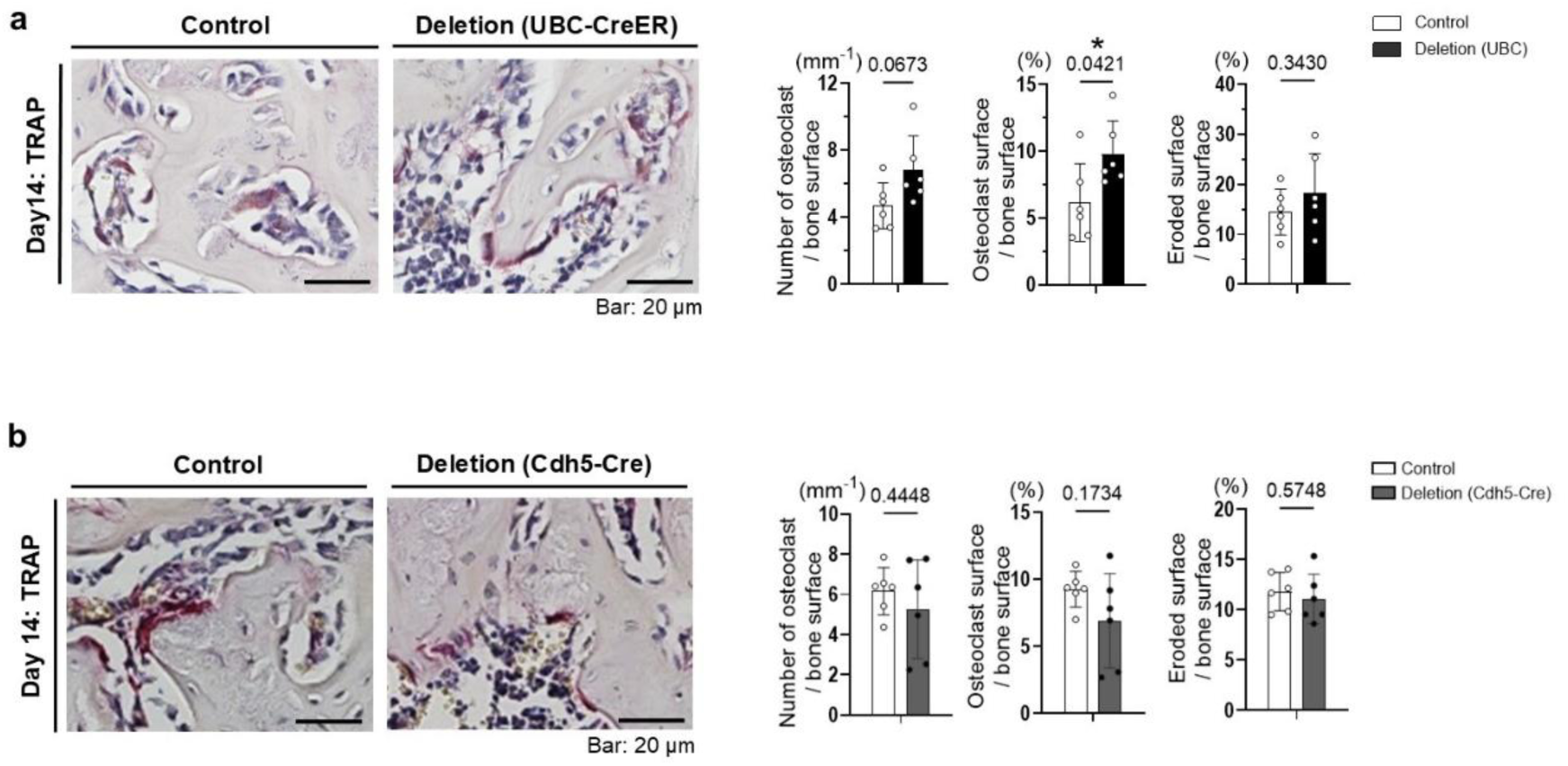
Osteoclast histomorphometric parameters after global and endothelial *Cx3cl1* deletion. a, Representative TRAP stained sections and quantification of osteoclast related histomorphometric parameters in the drill hole repair region of *Cx3cl1* flox/flox and UBC-CreERT2; *Cx3cl1* flox/flox mice at day 14 after femoral drill hole injury. Osteoclast number per bone surface, osteoclast surface per bone surface, and eroded surface per bone surface were quantified. n = 6 mice per group. b, Representative TRAP stained sections and quantification of osteoclast related histomorphometric parameters in the drill hole repair region of *Cx3cl1* flox/flox and *Cdh5*-Cre; *Cx3cl1* flox/flox mice at day 14 after femoral drill hole injury. Osteoclast number per bone surface, osteoclast surface per bone surface, and eroded surface per bone surface were quantified. n = 6 mice per group. Data are presented as mean ± SD. Each dot represents an individual mouse or independent sample. Statistical significance was determined using Student’s t test.

**Supplementary Fig. 12.**
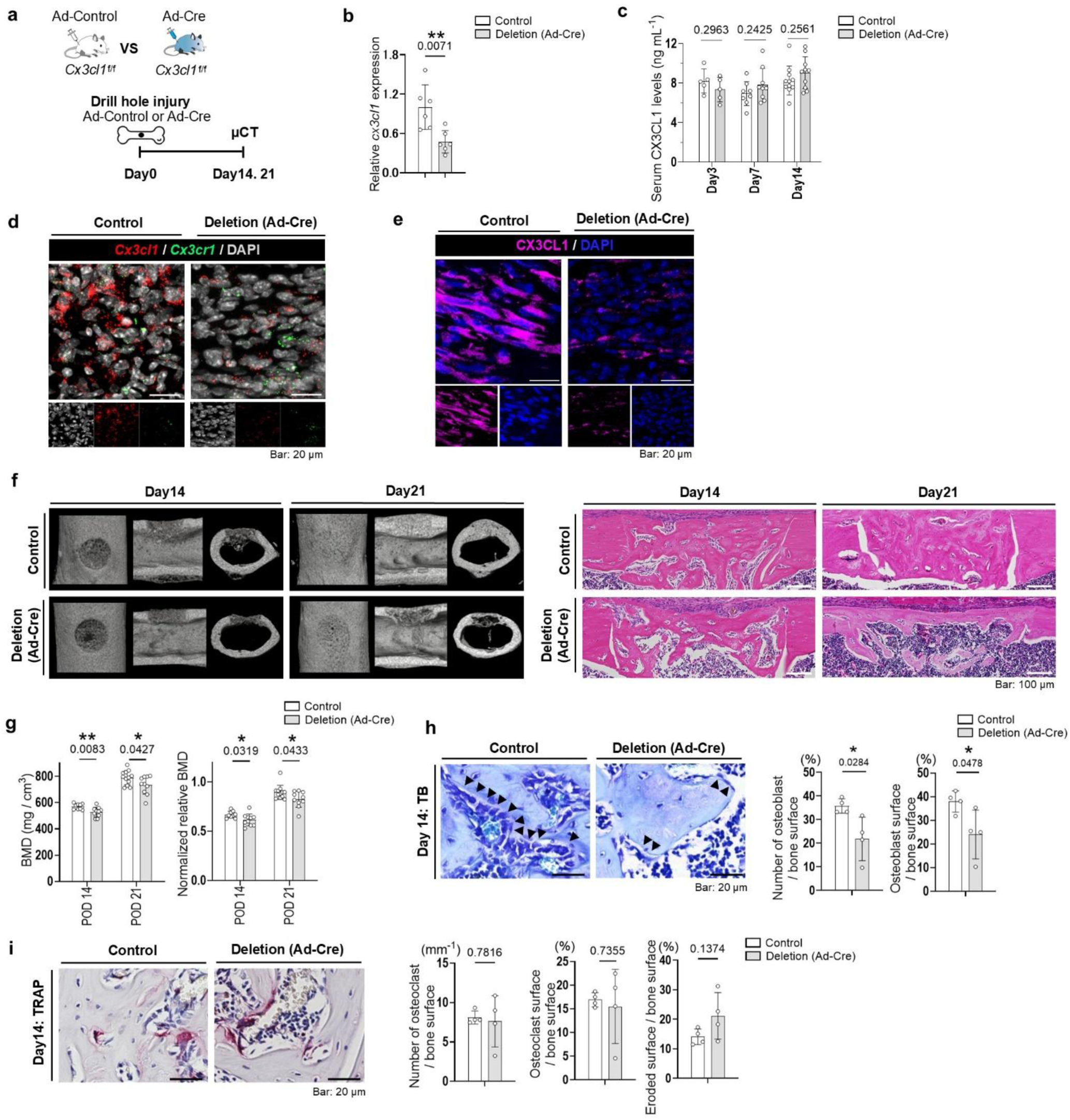
Local CX3CL1 deletion delays bone repair. a, Schematic experimental design for adenovirus mediated local deletion of *Cx3cl1* in young *Cx3cl1* flox/flox mice. Control adenovirus or Adeno-Cre was locally injected into the injury region, followed by femoral drill hole injury. b, *Cx3cl1* mRNA expression in injury adjacent muscle at day 3, normalized to *Gapdh*. *Cx3cl1* flox/flox mice treated with control adenovirus or Adeno-Cre. n = 6 mice per group. c, Serum CX3CL1 levels at days 3, 7, and 14 after injury. n = 5–11 per group. d, Representative RNAscope in situ hybridization images in the drill hole region of *Cx3cl1* flox/flox mice treated with control adenovirus or Adeno-Cre at day 7 after injury. n = 3 mice per group. e, Representative immunofluorescence images showing CX3CL1 (magenta) and DAPI (blue) in the drill hole region at day 7. n = 3 per group. f,g, Bone repair after local Cx3cl1 deletion. f, Representative μCT images and H&E sections at days 14 and 21. g, BMD and normalized BMD in the repair region. Normalized relative BMD was calculated as repair region BMD divided by contralateral cortical BMD. n = 9–10 per group at day 14 and n = 10–12 per group at day 21. h, Toluidine blue staining and osteoblast histomorphometry at day 14. Arrowheads indicate osteoblasts lining newly formed bone. Osteoblast number per bone surface and osteoblast surface per bone surface were quantified. n = 4 per group. i, TRAP staining and osteoclast histomorphometry at day 14. Osteoclast number per bone surface, osteoclast surface per bone surface, and eroded surface per bone surface were quantified. n = 4 per group. Data are presented as mean ± SD. Each dot represents an individual mouse or independent sample. Statistical significance was determined using Student’s t test.

**Supplementary Fig. 13.**
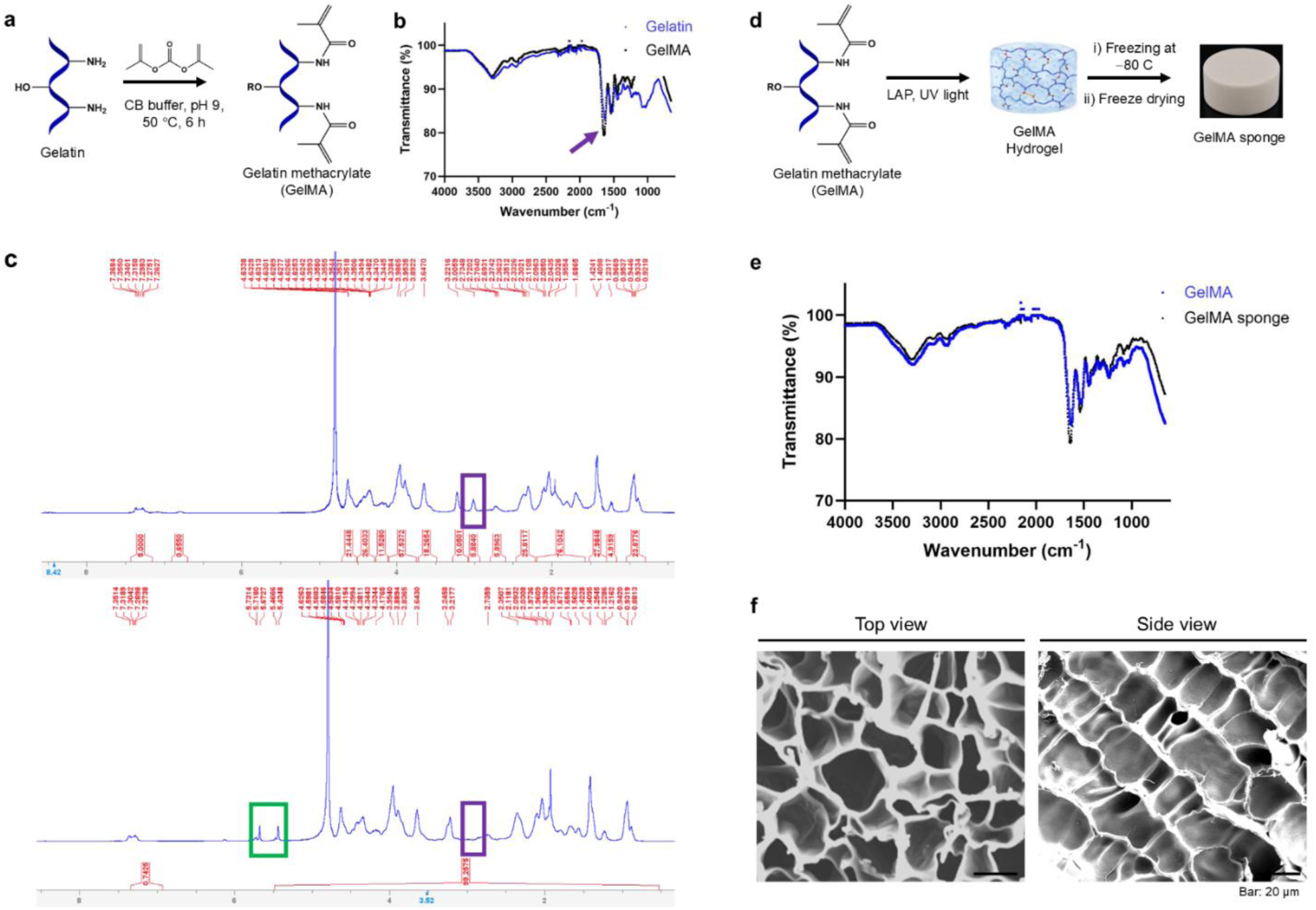
Synthesis and characterization of GelMA sponge. a, Schematic illustration showing the chemical modification of gelatin to GelMA via methacrylation. b, Fourier-transform infrared (FTIR) spectra of gelatin and GelMA acquired in attenuated total reflectance (ATR) mode using a germanium crystal. The arrow indicates the appearance of a peak at approximately 1640 cm⁻¹, consistent with C=C stretching vibration of methacrylamide groups in GelMA. c, ^1^H NMR spectra of gelatin (top) and GelMA (bottom) recorded on a 500 MHz instrument in D₂O. Successful methacrylation was confirmed by the appearance of methacrylate proton peaks at 5.46 and 5.71 ppm (green box). Extensive methacrylation of lysine free amine groups was further supported by a marked reduction in lysine ε-CH₂ proton signals. d, Schematic illustration showing the synthesis of GelMA sponge. Briefly, GelMA hydrogel was first prepared via photopolymerization of GelMA in the presence of the photoinitiator LAP under UV light. The resulting hydrogel was subsequently frozen at −80 °C and freeze-dried to obtain a porous GelMA sponge. e, FTIR spectra of GelMA precursor and GelMA sponge. f, Representative scanning electron microscopy (SEM) images of the freeze dried GelMA sponge showing its porous microarchitecture in top view and cross-sectional side view.

**Supplementary Table 1.**
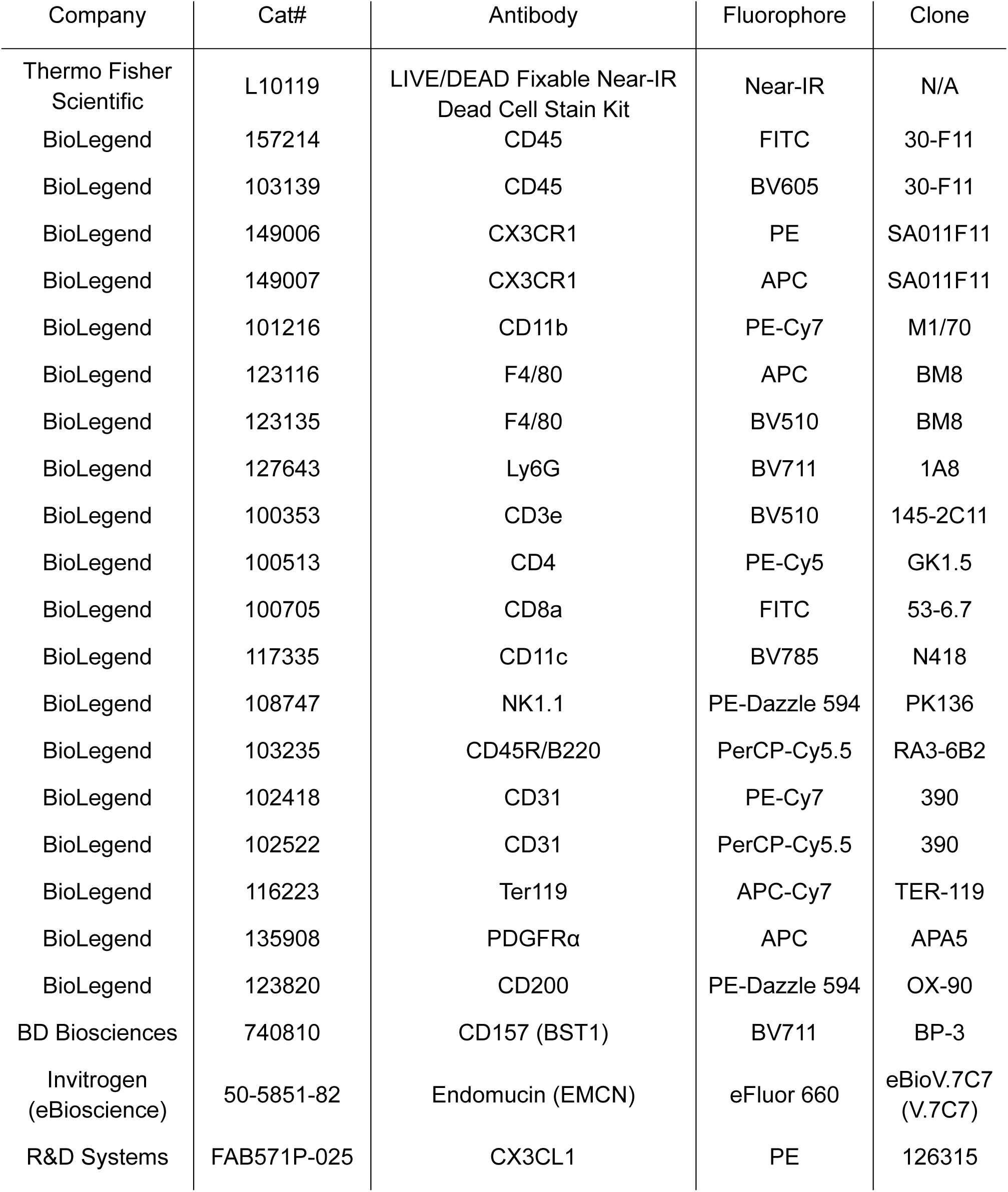
Antibodies used for flow cytometry.

